# A sensory system for mating in octopus

**DOI:** 10.1101/2025.10.28.685082

**Authors:** Pablo S. Villar, Hao Jiang, Tatiana Shugaeva, Emma L. Berdan, Arpita Kulkarni, Makoto Hiroi, Giovanni Masucci, Sam Reiter, Erik Lindahl, Rebecca J. Howard, Ryan E. Hibbs, Nicholas W. Bellono

**Author notes:** Correspondence should be addressed to R.E.H and N.W.B (,). These authors contributed equally to this work (Pablo S. Villar, Hao Jiang).

## Abstract

Sensory systems for mate recognition maintain species boundaries and influence diversification. Therefore, uncovering how molecules and receptors evolve to mediate this critical function is essential to understanding biodiversity. Male octopuses use a specialized arm called the hectocotylus to identify females and navigate their internal organs to reach the oviduct and deliver sperm. Here, we discovered that the hectocotylus is a dual sensory and mating organ that uses contact-dependent chemosensation of progesterone, a conserved ovarian hormone. We identify chemotactile receptors for progesterone and resolve the structural basis for their evolution from ancestral neurotransmitter receptors and subsequent expansion and tuning across cephalopods. These findings reveal principles by which sensory innovations shape reproductive behavior and suggest mechanisms for how sensory evolution contributes to the diversification of life.

## Introduction

Sensation is a gateway for reproduction across life by mediating the recognition of potential mates. Therefore, sensory receptors are evolutionary hotspots that can preserve conspecific recognition or drive interspecies mating, which underlies biodiversity (1–3). Thus, understanding how sensory receptors evolve and are tuned to recognize mating partners is critical to understanding the origins of new species.

Octopuses are solitary animals that mate upon infrequent encounters (4). Males use a specialized arm called the hectocotylus to identify females. The hectocotylus navigates toward and within the female’s mantle, distinguishing the oviduct and ovary from other organs to deposit a spermatophore that moves the length of the arm from the male mantle to the hectocotylus tip. Relatively few studies that are largely based on anecdotical evidence describe octopus mating, leaving important questions unanswered (5–11). How does the male recognize a correct mate during such rare encounters? How does the hectocotylus navigate the internal female organs to inseminate the oviducts?

Octopuses use their arms for ‘taste-by-touch’ chemotactile sensation of the seafloor, raising the possibility that mate recognition is associated with contact-dependent sensation (12). Octopus predation is driven by a complex, distributed arm nervous system (13–16) with chemotactile receptors (CRs) that diverged from ancestral neurotransmitter receptors, using structural adaptations to facilitate contact-dependent chemosensation of hydrophobic compounds (17, 18). While CRs have been shown to detect poorly soluble metabolites secreted by surface microbiomes of prey (19), it is unknown how octopuses detect mates.

Here, we report that octopus mating is driven by contact-dependent chemosensation in which males detect and mate with females in the absence of visual cues using the hectocotylus as a single organ for both mating and sensation. We discovered that the female cue for mating is the sex steroid progesterone, which is produced by conserved ovarian biosynthetic pathways. Progesterone activates male hectocotylus neural activity, autonomous movement, and is sufficient to elicit mating search behavior. Single-nuclei RNA sequencing demonstrates that the hectocotylus is enriched in sensory neurons and CRs and comparative approaches highlight the conservation of both signals and receptors across cephalopods. Physiology, histology, and cryo-EM structural biology reveal that the hectocotylus uses the chemotactile receptor for terpenes 1 (CRT1) as a high affinity progesterone receptor. Receptor elements that coordinate progesterone are rapidly evolving compared to conserved sites that are also used in prey detection. Thus, our study describes a novel sensory organ for mating and highlights how evolution can sculpt sensory receptors for mate detection, which is essential for the diversification of species.

## Results

### Octopus mating is driven by female chemosensory cues

To determine which sensory cues are required for octopus mating, we first separated two octopuses (wild-caught *Octopus bimaculoides*) by an opaque barrier with small openings through which they could introduce their arms in a controlled environment (**Fig. 1A**). To our surprise, even with minimal visual information, the male extended the hectocotylus through the barrier and carefully maneuvered toward and subsequently inserted the specialized appendage within the female mantle. Following insertion, the hectocotylus extended deep within the mantle and eventually stopped, at which point both the male and female paused all movement, sometimes for over an hour during spermatophore transfer (**Fig. S1A, B, Movie S1**). Similar results were obtained across numerous male-female pairs, even in the absence of visible light, but not when using male-male couples (**Fig. 1A**). While male-male couples interacted by touching their arms, they did not attempt mating, suggesting the existence of a female-specific cue (**Fig. 1A**).

**Figure 1.**
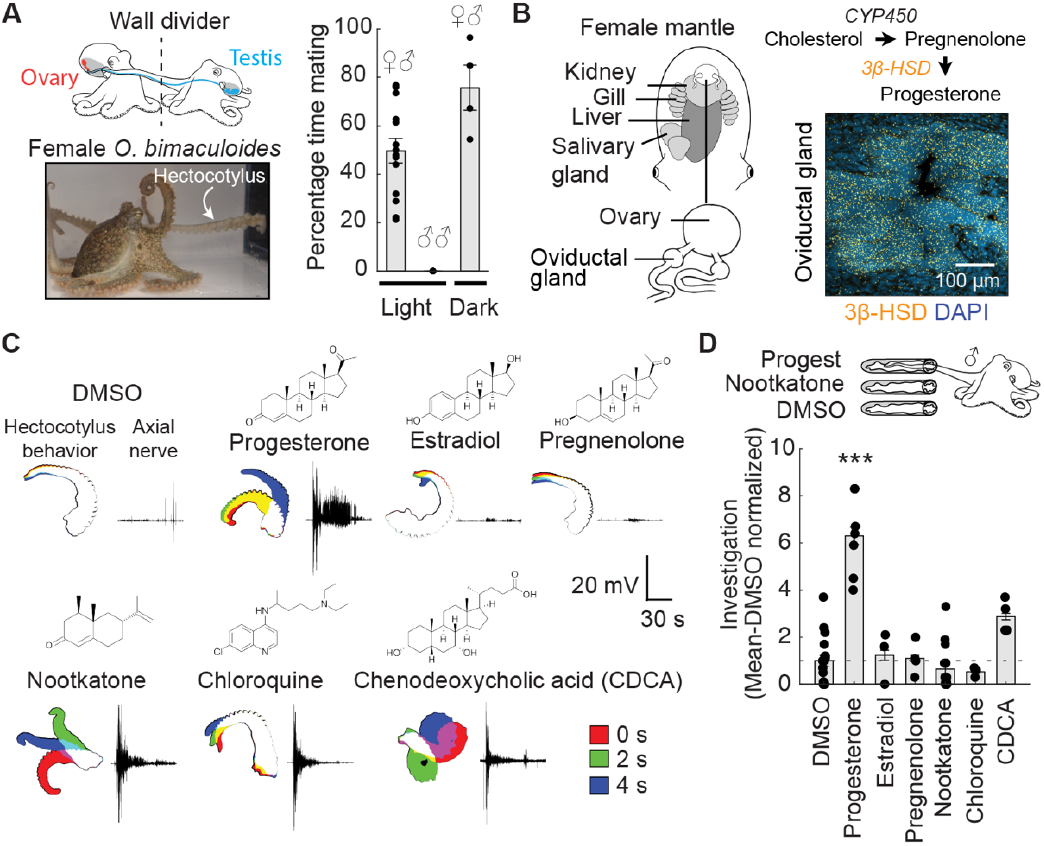
Octopuses use sex steroids to guide chemosensory behavior. **(A)** Left, *Octopus bimaculoides* mated without visual cues through a barrier with openings for the hectocotylus or in the absence of light. Right, couples spent majority of the time mating (defined as when the male hectocotylus reaches inside the female mantle) in both light (n= 3 mating pairs, 49.7 ± 5.1 %) and dark conditions (n= 2 couples, 75.7 ± 9.3 %). Male-male couples did not exhibit mating behavior, suggesting a female-specific signal (n= 3 couples). **(B)** Left, location of the ovary and oviducts in which sperm cells are deposited by the hectocotylus during mating. Right, the essential enzyme for progesterone biosynthesis 3β-hydroxysteroid dehydrogenase (3β-HSD, yellow) was expressed in the oviductal gland as visualized by in situ hybridization. Nuclei were stained with DAPI (blue). Representative of n= 3. **(C)** Amputated octopus hectocotyluses exhibited axial nerve and autonomous behavioral responses to steroids, bile acids and bitter molecules. Progesterone (100 *µ*M), but not the steroid precursor pregnenolone, elicited strong neural and behavioral responses (100 *µ*M, n= 3-6, p< 0.005, one-way ANOVA with Kruskal-Wallis test, summary data in Fig. S1F. **(D)** In mating behavioral trials, females were removed from the barrier tank and replaced with tubes in the openings for the hectocotylus. Male octopuses investigated tubes coated with progesterone more than vehicle, but did not explore structurally similar steroids, bile acids, or terpenes (n= 4–15 trials, data normalized to vehicle, p< 0.001, one-way ANOVA with Kruskal-Wallis test).

We next reasoned that a putative female-specific cue could originate from female-specific reproductive organs, namely the oviducts and ovary. Comparative tissue-based transcriptomics and gene ontology analysis showed a significant enrichment of biosynthetic enzymes important for the production of sex steroids and other known bioactive lipid molecules in the ovary and skin samples, consistent with observed steroid production during octopus mating season (20, 21) and conserved roles in animal reproduction and development (22, 23) (**Fig. S1C**). In situ hybridization confirmed that mRNA encoding the biosynthetic enzyme 3βhydroxysteroid dehydrogenase (3β-HSD), which is responsible for the conversion of the precursor pregnenolone into progesterone (**Fig. 1B**), was localized to the oviduct.

Considering that we observed markers for steroid biosynthesis in the oviducts, which are targeted by the hectocotylus during mating, we wondered whether the hectocotylus also detects sex steroids to guide mating behavior. We exploited the fact that severed arms exhibit neural activity and behavior to find that exogenously applied progesterone evoked robust hectocotylus activity (**Fig. 1C**). In contrast, structurally similar hormones like estradiol or the precursor pregnenolone failed to elicit such behavior (Fig. 1C). Indeed, the robustness of the response of the hectocotylus to progesterone was comparable to that observed for non-sexual arm signals such as terpenes, bitter molecules, bile acids, and touch (Fig. S1D–F). These results demonstrate that progesterone elicits a selective response that is locally detected and processed to trigger hectocotylus behavior.

To ask whether progesterone is sufficient to drive mating, we turned back to whole animal behavior with the same paradigm that we had used to define the chemosensory drive for mating. Similar to our initial experiments, the male and female were separated by a barrier, and the male searched for the female. Next, before mating took place, the female was replaced by conical tubes that were coated with chemical stimuli and attached to the small holes in the barrier in which the male inserted the hectocotylus to initiate mating. Strikingly, tubes coated with progesterone were sufficient to elicit mating search behavior with the male actively exploring the progesterone tube in place of searching the female mantle (**Fig. 1D, Movie S2**). This contrasted with tubes coated in other steroid derivates, bile acids, terpenes, or bitter molecules, which were previously shown to elicit aversive behavior in predation and exploration by normal arms (**Fig. 1D**). Thus, we concluded that progesterone is a robust signal for octopus mating

### The hectocotylus is a conserved sensory and mating organ

While normal arms are used for chemotactile exploration and predation, the hectocotylus is almost exclusively used for mating and often even protected during hunting (4, 24). During copulation, sperm packages are transferred from the testis to the base of the hectocotylus and slid down the arm through an external skin groove to the arm tip for release near the oviducts (**Fig. 2A**). Is the hectocotylus also a specialized sensory organ? A clue that this might be the case came from scanning electron microscopy, which revealed that the hectocotylus tip was covered by small sucker cups, like those found in regular arms and used in chemotactile sensation of the seafloor and prey (Fig. 2B). Furthermore, tissue clearing and immunohistochemical staining revealed dense neuronal innervation of the hectocotylus sucker cups, comparable to a normal arm tip (**Fig. 2C**). These anatomical observations are consistent with a direct sensory role for the hectocotylus, as suggested by the physiology and behavior of isolated arms (**Fig. 1C, S1D–F**).

**Figure 2.**
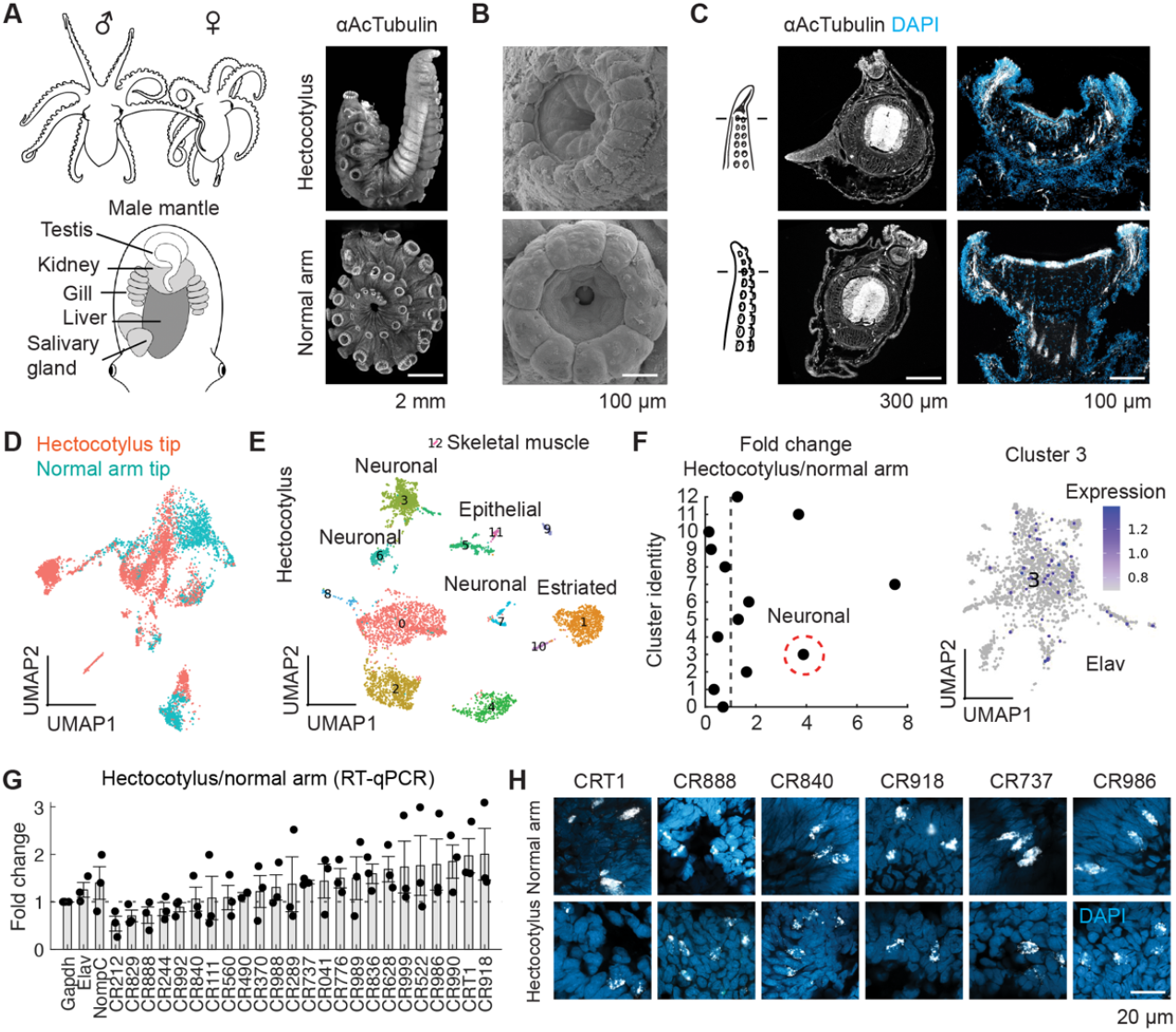
The hectocotylus is a chemosensory and mating organ. **(A)** Left, spermatophores are generated in the mantle, transported down the hectocotylus arm to the tip, and deposited at the oviduct openings in the female mantle. Right, tissue-cleared arm tips stained with the neuronal marker acetylated α-tubulin. **(B)** Scanning electron micrographs showed sucker cups in the hectocotylus tip were similar to the normal male arm. **(C)** Cross-sections of hectocotylus (top) and normal arm (bottom) immunostained against α-acetylated tubulin (white). DAPI nuclear staining was used to visualize the overall tissue morphology. Representative of n= 3. **(D)** UMAP reduction of single nuclei RNA sequencing (snRNA-seq) data from sucker sensory epithelium in the hectocotylus tip (4266 nuclei) versus the normal arm tip (2702 nuclei) showed clear differences in cellular populations, while the arm base was the same (see also Fig. S2). **(E)** UMAP of clustered hectocotylus cells shows multiple neuronal clusters, which were identified by the expression of the neuronal marker gene Elav in at least 10% of cells. **(F)** The largest neuronal cluster (#3) represents 25% of all hectocotylus tip cells (1067/4266) versus 6.4% in the normal arm (174/2702). **(G)** Quantitative RT-PCR demonstrated that the hectocotylus expresses chemotactile receptors (CRs), with many at higher levels than the normal arm (n= 3 animals, ranked and normalized to the housekeeping gene Gapdh). Dotted line indicates equal expression in hectocotylus and normal arm. **(H)** CRs localize to the hectocotylus tip sensory epithelium as shown by in situ hybridization (white staining). Nuclei were stained with DAPI (blue). Representative of n= 6 animals.

To ask whether the hectocotylus exhibited molecular and cellular features indicative of a sensory organ, we used comparative single nucleus RNA sequencing (snRNA-seq) to profile the hectocotylus tip that contacts the ovaries versus the base arm, which is similar to the other nonspecialized arms used for predation. Following profiling of ∼13,000 nuclei across arms, we classified cellular clusters based on enriched transcripts and found that the base arms contained similar cellular populations as defined by a combination of cell types and differential expression (**Fig S2A**). In contrast, profiling revealed three distinct populations that were expanded in the hectocotylus versus normal arm tips (**Fig. 2D–F**). Two hectocotylus-enriched clusters were related to metabolism and secretion, and the third (Cluster 3) was neuronal and enriched for the gene ontology (GO) term “response to chemical stimulus” (**Fig. S2B–G**). Although sensory receptors are often expressed at low abundance, even snRNA-seq showed that Cluster 3 contained CRs, the mechanoreceptor NOMPC, and the neuronal marker Elav, collectively suggesting a sensory function (**Fig. S2C**). Quantitative RT-PCR and in situ hybridization confirmed that CRs are robustly expressed in the epithelium of the small suckers found along the hectocotylus tip (Fig. 2G, H). Thus, the hectocotylus exhibits all the molecular and cellular hallmarks of a multimodal sensory organ for chemosensation and mechanosensation, consistent with our physiological and behavioral data.

Because sensory systems for mating are central to evolutionary fitness, we hypothesized key anatomical and molecular components would be present across cephalopods. To test this possibility, we compared the putative sensory systems for mating in *Octopus bimaculoides* with phylogenetically and geographically diverse octopods (*Octopus rubescens* and *Abdopus aculeatus*) and a decapod (*Euprymna berryi*) (**Fig. 3A**). First, we found that ovaries from all cephalopods expressed conserved biosynthetic enzymes for producing sex steroids, including progesterone (Fig. S3A). Similarly, the hectocotylus across all species contained anatomically similar sucker cups (Fig. 3B) and robustly responded to exogenously applied progesterone and other sex steroids to a varying extent across species (Fig. 3C, Fig. S3B). Responses to progesterone were comparable to CR ligands sensed by nonsexual arms, such as terpenes, bitter molecules, and bile acids (**Fig. 3C, Fig. S3B**). Importantly, the molecular markers for sensory function were conserved as CRs were enriched in the hectocotylus tips of all species (**Fig. 3D, Fig. S3C**). Finally, consistent with these findings, the phylogenetically distant *Abdopus aculeatus* also mated in the absence of any visual cue (**Fig. S3D**), suggesting that chemosensory driven mating is not limited to *Octopus bimaculoides*.

**Figure 3.**
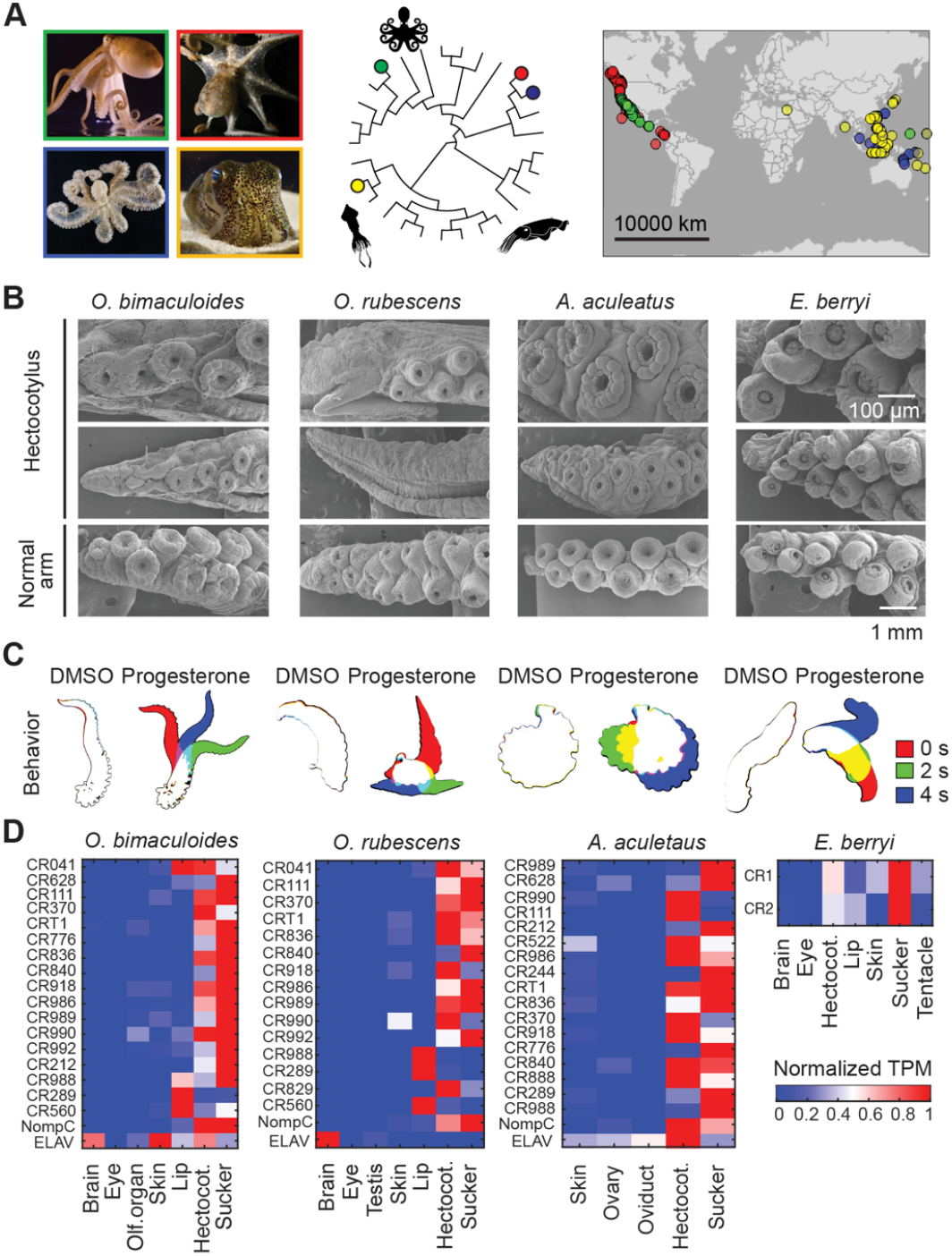
Steroid sensing is conserved among cephalopods. **(A)** Octopus and squid species analyzed for behavior and transcriptomics with their phylogeny and geographic distribution: *Octopus bimaculoides* (green), *Octopus rubescens* (red), *Abdopus aculeatus* (blue) and *Euprymna berryi* (yellow). **(B)** Comparative ultrastructural analysis of hectocotylus arm tips across cephalopod species. Scanning electron micrographs for normal and hectocotylized arms are shown for *O. bimaculoides, O. rubescens, A. aculeatus and E. berryi*. Representative of n= 4. **(C)** Hectocotyluses from all tested species were sensitive to the sex steroid progesterone versus vehicle (*O. bimaculoides*, n= 3; *O. rubescens*, n= 4; *A. aculeatus*, n= 8; *E. berryi*, n= 7, p < 0.004). See summary data for other steroids, bile acids and terpenoids in Fig. S3B. **(D)** CR transcripts were enriched in the hectocotylus of *O. bimaculoides, O. rubescens, A. aculeatus and E. berryi* compared to other sensory tissues. The color scale shows transcripts per million (TPM) normalized across samples.

### CRT1 is a rapidly evolving progesterone receptor in octopus

We next sought to identify the progesterone receptor of the hectocotylus. We started by screening a panel of 800 bioactive lipids, including steroids, across five different CRs that were most abundantly expressed in the hectocotylus and could be functionally probed in expression systems (**Fig. S4A–C, Table S1**). We found that only one CR, CRT1, robustly responded to sex steroids, including progesterone (**Fig. 4A, B, Table S2**). Patch-clamp electrophysiology with CRT1 demonstrated that progesterone is the highest affinity natural ligand for an octopus CR reported to date (**Fig. 4C**, (19)). This response was also selective because CRT1 was insensitive to the structurally similar precursor pregnenolone, matching the pharmacological profile of hectocotylus neural activity and behavior (**Fig. 4C**). Indeed, cryo-EM structural analysis of CRT1 bound to progesterone demonstrated that progesterone was coordinated within the canonical orthosteric binding site (**Fig. 4D-E, Fig. S5A**). The density for progesterone was strong but ambiguous as to which orientation the steroid was bound in the pocket. We used molecular dynamics (MD) simulations to identify the most energetically favorable orientation of progesterone in the cryo-EM density map (**Fig. S5B, C**). Substitution of residues coordinating progesterone drastically reduced progesterone-evoked activity (**Fig. 4F**), supporting the inference that this solvent-exposed site being is the route through which the lipidic agonist acts.

**Figure 4.**
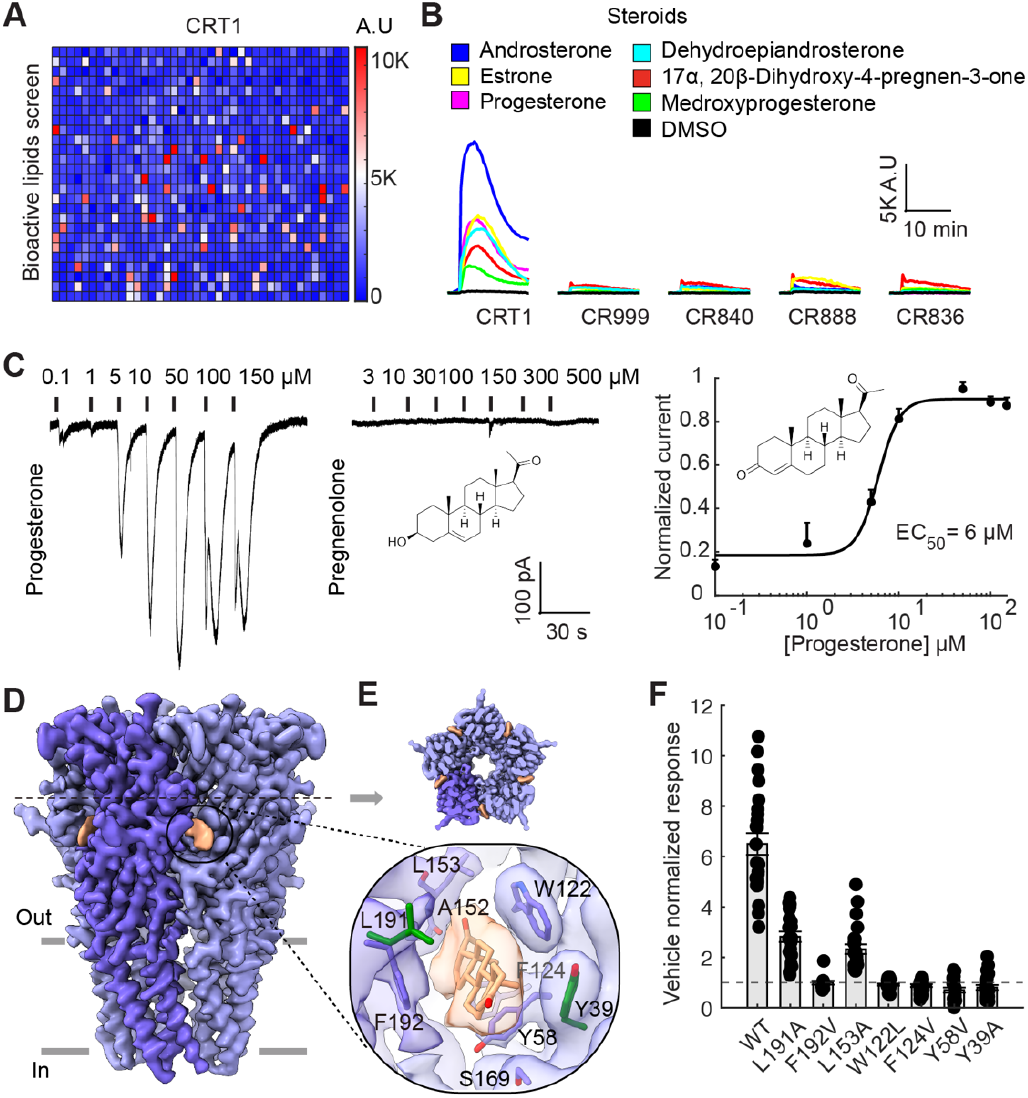
CRT1 is a steroid receptor with high affinity for progesterone. **(A)** Calcium responses of HEK293 cells expressing *O. bimaculoides* CRT1 to a panel of 800 bioactive lipid molecules (10 *µ*M) identifies progesterone as a robust candidate ligand. Responses were quantified by the increase in fluorescence of GCaMP6 expressing cells as an indicator of CR activity over basal. Each cell represents the top response to a single molecule. Color bar indicates fluorescence in arbitrary units. **(B)** Comparison across all tested CRs highlights CRT1 sensitivity to sex steroids. Traces represent raw responses from single trials shown in panel A. **(C)** Whole-cell currents from HEK293 expressing CRT1 responded to progesterone but not the precursor pregnenolone (n = 6, EC50 = 5.8 ± 0.5 *µ*M). **(D)** Cryo-EM density map of octopus CRT1 bound to progesterone. Progesterone is highlighted in orange. **(E)** Upper, top view of the cryo-EM map. Lower, map and atomic model of the orthosteric ligand binding site showing coordination of the most favorable progesterone pose. Interacting residues are shown as sticks. **(F)** Point mutations of residues coordinating progesterone abolished responses compared to wild type CRT1. Bar plot with vehicle normalized peak responses for CRT1 wildtype and mutants in an *in vitro* calcium assay (n= 8–24, p< 0.001, one-way ANOVA with Kruskal-Wallis test).

CRT1 was previously characterized as a receptor for metabolites produced by the microbiota on surface of prey (**Fig. 5A**) (19). How does the same receptor mediate both steroid recognition for mating and metabolite sensing for predation? By comparing the structures of CRT1 bound to norharmane (predation) or progesterone (mating), we found that both had an open pore conformation and a similar permeation pathway, consistent with their ability to activate CRT1 (Fig. S6). Yet, progesterone had an increased interaction area and inferred binding energy with three residues of the orthosteric binding site compared to that for norharmane, which are: L153, L191, and Y39 (**Fig. 5B, E, Fig. S7A**).

**Figure 5.**
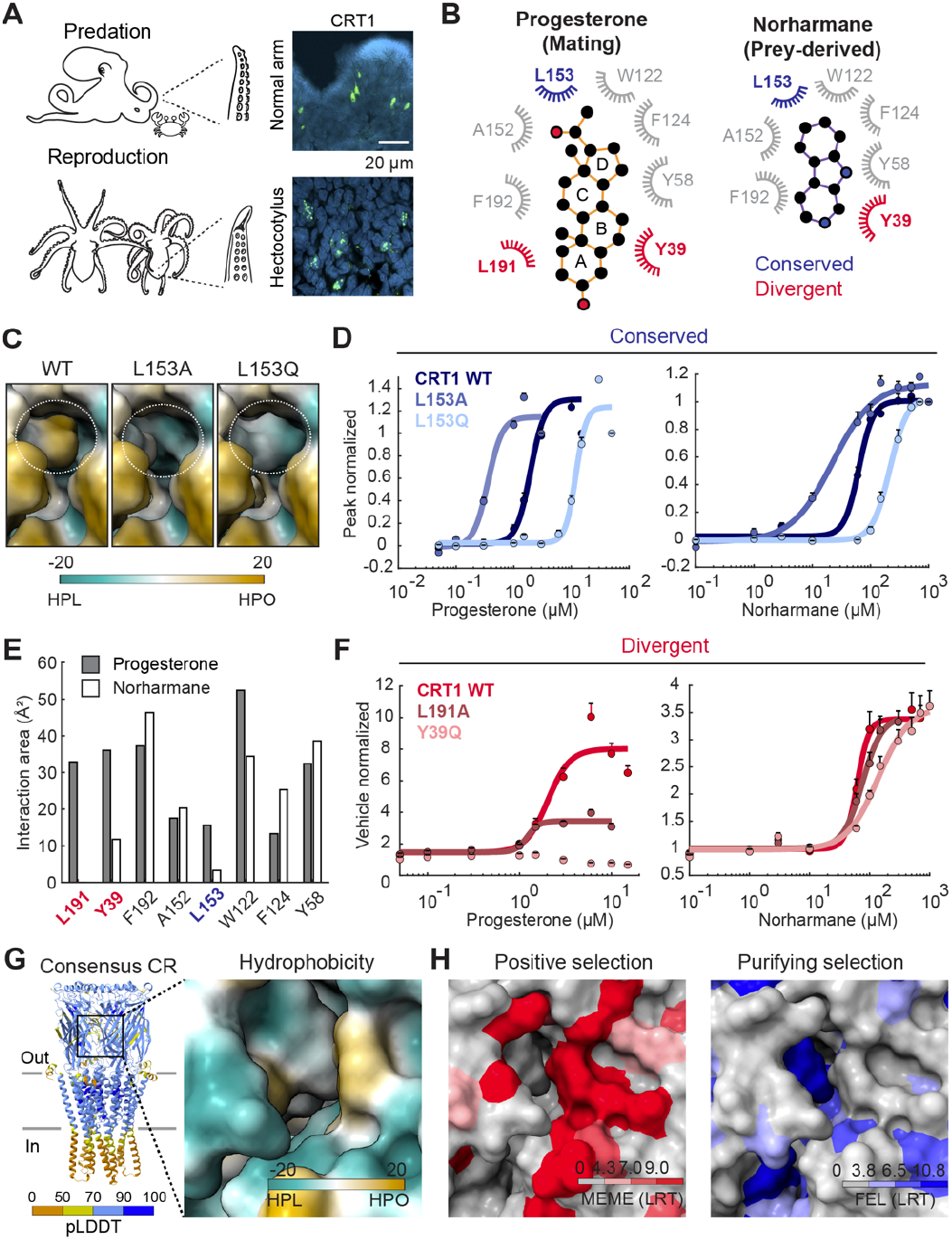
Structural adaptations of CRT1 underlie high specificity to steroids. **(A)** CRT1 expression in the normal arm and hectocotylus of octopuses support predation and mating behaviors. CRT1 localizes to the normal arm and hectocotylus tip sensory epithelium as shown by in situ hybridization (green staining). Nuclei were stained with DAPI (blue). Representative of n= 8 **(B)** Binding pocket residues coordinating progesterone (left) and norharmane (right) highlighting the conserved residue L153 colored in blue and divergent residues L191 and Y39 colored in red. Structural and selection analyses in Fig. S7. **(C)** Pocket comparison among WT, L153A and L153Q. L153A exhibits a larger and deeper binding pocket compared to the WT, whereas L153Q shows a comparable volume but increased hydrophilicity. Dashed white circles indicate L153 position. **(D)** Mutating the conserved L153 into a smaller residue (L153A) increases binding affinity for progesterone (left) and norharmane (right) while a larger residue (L153Q) decreases CRT1 sensitivity to both molecules. Progesterone EC50 WT= 2.2 ± 0.2 *µ*M, n= 24; L153A= 0.4 ± 0.0 *µ*M *µ*M, n= 24; L153Q= 12.3 ± 0.5 *µ*M, n= 16. Norharmane EC50 WT= 75.6 ± 6.6 *µ*M, n= 32; L153A= 19.8 ± 2.0 *µ*M, n= 24; L153Q= 247.7 ± 19.4 *µ*M, n= 16. **(E)** Interaction area of residues L191, Y39 and L153 with progesterone is higher compared to norharmane. **(F)** Mutating residues under positive selection L191 and Y39 affects progesterone binding but not norharmane. Mean progesterone peak response (15 *µ*M, vehicle normalized) WT= 6.5 ± 0.4, n= 24; L191A= 2.8 ± 0.2, n= 24; Y39Q= no response, n= 16. Norharmane response (150 *µ*M, vehicle normalized) WT= 2.6 ± 0.2, n= 28; L191A= 2.8 ± 0.2, n= 16; Y39Q= 2.0 ± 0.2, n= 12. **(G)** Predicted AlphaFold3 (28) structure of a consensus CR. A consensus protein sequence was obtained from 235 CR sequences across 29 cephalopod species including octopus, squid and cuttlefish species. Right, hydrophobicity of the predicted ligand binding pocket in a cephalopod consensus CR. Superposition of the structures shows that progesterone from the cryo-EM structure fits the predicted pocket. **(H)** Residues in the binding pocket colored by the likelihood ratio test (LRT) value for the positive selection pressure test MEME (left) and purifying selection test FEL (right).

The conserved residue L153 forms the back wall of the conserved orthosteric pocket, coordinating the acetyl chain on the D-ring of progesterone (**Fig. 5B, Fig. S7B**). We reasoned that if this conserved region provides a general groove for hydrophobic ligands, then altering its hydrophobicity should similarly affect binding of both progesterone and norharmane. Indeed, substituting L153 for a hydrophobic alanine residue increased binding affinity for both ligands, while the hydrophilic residues (glutamine and asparagine) decreased affinity (**Fig. 5C, D, Fig. S8A–E**). Additionally, replacing other conserved residues within the hydrophobic groove significantly reduced binding of both ligands (**Fig. S7D**). These results suggest a conserved site for coordinating poorly soluble ligands important in contactdependent aquatic sensation of predatory metabolites or steroids in mating.

In contrast, L191 and Y39 formed a distinct hydrophobic sandwich around the A-ring at the bottom of progesterone with increased interaction area compared to norharmane, which extended beyond the canonical binding pocket for norharmane (**Fig. 5B, E**). Intriguingly, these residues were under diversifying selection, indicating that they are rapidly evolving versus the other residues in the conserved pocket that coordinates both molecules (**Fig.S7C**). Consistent with these observations, mutating L191 and Y39 showed a stronger effect on progesterone sensitivity when compared to norharmane (**Fig. 5F, Fig. S8A–E**). Collectively, these results suggest that CRT1 has functionally diversified in two ways: 1) CRT1 is expressed in sensory suckers of both the normal arm and mating hectocotylus; and 2) it has evolved a hydrophobic binding pocket that loosely binds molecules important for predation to a high affinity steroid binding pocket for molecules involved in mating (**Fig. 5A**).

While the hectocotylus from all cephalopods was sensitive to progesterone, we noticed varying sensitivities to related steroid derivates (**Fig. S3B**). As such, we wondered whether CRs are broadly diversifying structural regions that could tune steroid binding to support species-specific chemosensory behaviors, including recognizing mating partners. To ask this question, we used genomic sequences from 30 cephalopod species (235 total receptors) to build a consensus receptor and used AlphaFold3 to predict its structure. The consensus orthosteric site resembled the relatively flat hydrophobic pocket of CRT1, indicating a common biophysical property of CRs that allows binding of relatively insoluble molecules (Fig. 5G). We then overlaid the likelihood ratio test (LRT) value for positive (MEME) and purifying selection (FEL) to determine which regions are under evolutionary pressure (Fig. 5H). These analyses revealed conservation of the inner pocket, the same region important for both progesterone and norharmane sensitivity in CRT1, and diversifying selection at the outer region that mediates high affinity for progesterone. This contrasted with ancestral nicotinic acetylcholine receptors, which exhibited generally conserved binding sites compared with more recently derived CRs (Fig. S7E–G). These results suggest that CRs evolved a core binding motif suited for “taste by touch” chemotactile sensation of poorly soluble molecules in aquatic environments. This function underwent diversification, enabling distinct behaviors ranging from predation to mating by evolution of the outer binding pocket.

## Discussion

Chemoreceptor families expand and contract to meet lineage-specific ecological and behavioral demands. Like vomeronasal and variant ionotropic receptors in other animals, CRs evolved from an ancient neurotransmitter receptor scaffold. In contrast to animals that evolved distinct receptor families to support diverse behaviors, octopuses employ a single family for predation, maternal behavior, and mating. In a wide range of animals, chemosensitive and reproductive organs remain separate: pheromones are detected by dedicated sensory receptors and epithelia, such as in the rodent vomeronasal organ (25), whereas mating organs act as effectors. The octopus hectocotylus breaks this modularity by collapsing chemosensation and sperm transfer into a single appendage, similar to the male reproductive organ of some insects (26, 27). This integrated function ensures mate assessment at the site and moment of gamete delivery, a strategy well suited to the rare encounters of these solitary animals.

Cephalopods represent a rich source of biological novelty. Along with abandoning the protection afforded by the ancestral molluscan shell, these animals evolved longdistance locomotion, active predation, and problem solving. This shift coincided with the evolution of flexible arms, an advanced distributed nervous system, and CRs as a novel sensory receptor expansion that underpins their ecological success. Among the most profound departures from their molluscan relatives is reproduction. Whereas bivalves and gastropods rely on hermaphroditism and broadcast external spawning, cephalopods evolved separate sexes, sex chromosomes, and direct transfer of spermatophores. The evolution of sexual reproduction between distinct male and female imposes selective pressure on precise mate recognition and timing. Thus, by merging precise sperm transfer and sensory novelties suited to contact-dependent chemosensation, the cephalopods hectocotylus may have reinforced species-specific sensory boundaries, contributing to reproductive isolation and facilitating diversification.

Across animals, sensory innovation and reproduction are repeatedly entwined. Insects use refined pheromone receptors for species recognition, mammals diversify sensory repertoires of receptors involved in mate choice, and cephalopods remodel arms into combined chemosensory– reproductive organs. In each lineage, evolution has reshaped conserved receptor scaffolds to serve reproductive systems, and cephalopods push this integration further by fusing sensation and sperm transfer into a single appendage. This fusion not only secures reproductive efficiency but may have generated new pathways that contribute to lineage diversification.

## Supporting information

Movie S1. Octopus mating

Movie S2. Tube assay

## Acknowledgements

We thank C. Winkler and B. Grasse for providing octopuses and squids; B. Walsh and P. Kilian for assistance with animal care; A. Grearson for photography and illustrations; C. Saunders for assistance with MD simulations; The Harvard Electron Microcopy facility; The Harvard Center for Biological Imaging (HCBI); The Bauer Core facility; The UC San Diego Cryo-EM Facility. This research was supported by NIH grant R24GM154185 and performed at the Pacific Northwest Center for Cryo-EM (PNCC) with assistance from Omar Davulcu, and NIH grant R01NS129060 to NWB and REH. PSV was supported by the Jane Coffin Childs Memorial Fund for Medical Research and HJ by the Kavli Institute of Brain and Mind.

## Author contributions

PSV and NWB conceptualized the study. PSV, HJ, TS, ELB, AK, MH and GM contributed to molecular, cellular, structural and behavioral experiments. All authors were involved with writing or reviewing the manuscript.

## Data and materials availability

Sequencing data generated from this study has been deposited to the BioSample database under the accession number PRJNA1336213. The atomic model and cryo-EM map of CRT1-progesterone have been deposited in the Protein Data Bank and Electron Microscopy Data Bank under the IDs PDB 9YI4 and EMDB EMD-72977, respectively.

## Materials and Methods

### Animals

All experiments were conducted following the guidelines of the Harvard Animal Care and Use Committee. Adult male and female California two-spot octopuses (*O. bimaculoides*), ruby octopuses (*O. rubescens*) and abdopuses (*A. aculeatus*) were wild caught (Aquatic Research Consultants) while adult squids (*E. berryi*) were bred at the Marine Biological Laboratory (Marine resource Center, Woods Hole, MA, USA). *O. bimaculoides* and *O. rubescens* were kept in filtered natural sea water at 20 °C and 17 °C, respectively, at pH 8, salinity of 32 ppt and under a 12h light/dark cycle inside 23 L tanks connected to a recirculating closed system (320 L). Abdopuses were kept in filtered natural seawater at 25 °C, pH 8 and salinity of 35 ppt inside 14 L tanks connected to a recirculating closed system (1500 L) and under a 12 h light/dark cycle.

Octopuses were daily fed with fiddler crabs (Northeast Brine Shrimp, Oak Hill, FL) and squids fed with grass shrimp (Aquatic Indicators). All animals were anesthetized using a standardized protocol with sedation in 3% ethanol in sea water followed by decapitation.

### Cells

HEK293T cells (ATCC, authenticated and validated as negative for mycoplasma by the vendor) were grown in DMEM, 10% fetal bovine serum (PEAK Serum) and 50 IU/mL penicillin and 50 µg/mL streptomycin (Gibco) at 37 °C and 5% CO2 using standard techniques. Cells were transfected in Opti-MEM Reduced Serum Media (Gibco) with Lipofectamine 2000 (Invitrogen) for 6h at 37°C and following the manufacturer protocol. 1 µg of CR plasmid and 0.5 µg of GCaMP6s DNA were cotransfected for Ca2+-based experiments while 0.3 µg of GPF was cotransfected for whole cell recordings. After transfection, Opti-MEM was aspirated and replaced by DMEM, and cells were let to express the plasmids overnight at 37°C. Before experiments, DMEM was replaced by mammalian Ringer’s solution (see below). Gene cloning, synthesis and mutagenesis was performed by GenScript. Plasmids (Table S3) were amplified by transformation of Stellar competent cells (TaKaRa) and liquid culture followed by purification using a Maxiprep kit (ZymoPURE).

### Mating behavior

For octopus behavior, the mating arena consisted of an acrylic container of 60×20×25 cm (W, L and H) with an opaque black acrylic divider with three small openings (8 mm in diameter) at the bottom. Male and female octopuses were transferred to the behavior tank filled with natural sea water and let freely swim and investigate under light or dark conditions. Male or female octopuses self-initiated each mating trial by reaching through the wall openings. Trials were performed in consecutive days and for at least 1h or until the end of the copulatory behavior and voluntary separation. The copulatory behavior was often repeated during the same trial and rarely not observed. Experiments were recorded with top and side view cameras (GoPro) under white light or in the dark using an FLIR camera (Teledyne) under infra-red illumination. Mating behavior was classified as precopulatory (swimming and exploring) and copulatory when the hectocotylus was introduced inside of the conspecific mantle. After each behavioral trial, octopuses were transferred to their home tanks. For abdopus behavior, animals were wild caught during night low tide on tidal flats, sexed by identifying the hectocotylus, moved to a mating arena and kept separated with a barrier containing 16 holes (6 mm diameter each). Filming sessions were conducted both under light and in the dark. Filming was conducted over 27 h per mating pair.

### Mating tube behavior

During a few catch trials, male octopuses previously exposed to females were transferred to the mating behavior tank which had three tubes in the place of the wall openings. The inside wall of the tubes was covered by 2% agar containing the test molecules or vehicle control and octopuses were let to investigate with their arms. Octopuses self-initiated each trial by reaching through the wall while recorded with a top camera (GoPro). The total time spent investigating each tube was quantified and normalized to the mean investigation time of the control tube (DMSO).

### Semi-autonomous arm behavior

Arm tips were placed in a chamber containing filtered natural sea water and held by suction at the center of the field of view. Chemicals were locally applied in volumes of 200 µL and at a concentration of 100 µM diluted in natural sea water. The behavior of the arm was recorded from the top with a FLIR camera (Teledyne) at 30Hz. Videos were down sampled, filtered with a gaussian blur, thresholded and binarized to extract the overall arm features (ImageJ). Overall arm movement was quantified using the kymograph plugin applied across frames.

### *In vitro* calcium assay

Cells were cultured in clear 96-well plates and cotransfected with specific CR genes and GCaMP6s overnight as described above. The culture media was replaced by mammalian Ringer’s solution (see below) and cells kept at room temperature for at least 1h for signal to reach low baseline levels. Fluorescence measurements on GCaMP/CR expressing cells were performed using the BioTek Synergy Neo2 plate reader (Agilent) from total induced fluorescence emitted from the top. For dose-response curves, chemicals were prepared at 5x dilutions in a v-bottom 96-cell plate and added to the cells after 3 min of baseline measurements using a 96-channel pipette (MicroPro300, RAININ). For small molecules screens, we used the Bioactive lipid I screening library (Cayman) prepared at 3x dilutions from 1 mM stocks in DMSO and added to the wells with a 96-channel pipette. Fluorescence in each well was measured every 20 s and responses quantified as the peak of the calcium trace after chemical application.

### Patch clamp electrophysiology

Whole-cell recordings were conducted from cultured cells in a perfusion chamber (Warner Instruments) using a MultiClamp 700B amplifier (Axon instruments). The signal was low pass filtered at 3 kHz and acquired at 10 kHz using the Digidata 1550B interface (Axon instruments). The amplifier was controlled using the pCLAMP software (Axon instruments). Cells were visualized using an inverted fluorescence microscope with DIC contrast (Olympus IX73). Whole-cell currents were recorded at a holding potential of –80 mV using pipettes filled with an internal solution of the following composition (in mM): 140 CsMeSO4, 5 NaCl, 1 MgCl2, 10 HEPES, 10 EGTA and 10 Sucrose. The pH of the internal solution was adjusted to 7.2 with CsOH and final osmolarity was 290 mOsm. Patch pipettes were pulled using a horizontal puller (P-97, Sutter Instrument) from thick-wall borosilicate glass capillaries (Sutter Instrument), having a resistance of 3–6 MOhm. A mammalian Ringer’s solution was used as extracellular solution and contained in mM: 140 NaCl, 5 KCl, 10 HEPES, 10 glucose, 2 CaCl2, 2 MgCl2. The pH of the ringer’s solution was adjusted to 7.2 with NaOH. Micro perfusion of chemicals was performed using the SmartSquirt Micro-Perfusion system (Automate Scientific) controlled by the pCLAMP software. Data were analyzed using custom MATLAB (MathWorks) software.

### Axial nerve recording

Arm tips were severed from deeply sedated animals and placed in a Petri dish containing filtered natural sea water. Differential extracellular recordings were performed with a bipolar suction electrode (A-M systems) on the open end of the axial nerve cord using a Warner DP-311A amplifier. The signal was band passed filtered between 10Hz and 10KHz and digitized using a Digidata 1440A interface (Axon instruments). The amplifier was controlled using the Clampex software (Molecular devices). Chemicals were locally applied in volumes of 100 µL and at a concentration of 100 µM diluted in natural sea water. Data were analyzed using custom MATLAB software.

### Transcriptomics

*De novo* transcriptomes were generated from tissues of *O. bimaculoides, O. rubescens, A. aculeatus* and *E. berryi*. All tissues were flash frozen in RNAlater (Sigma) and kept at –80°C until use. RNA extraction, library preparation and sequencing were performed by Genewiz (Azenta). Adapter and quality trimming was performed using Trim Galore and transcriptomes were assembled using Trinity. Candidate coding regions were identified using TransDecoder and transcript abundance quantified using Kallisto. Gene annotation was performed using Diamond. Sequencing data generated from this study are available at the NCBI Sequence Read Archive under the BioProject accession number PRJNA1336213.

### Immunohistochemistry

Tissue was fixed with 4% paraformaldehyde in PBS overnight at 4°C. Fixed tissue was embedded in OCT and frozen at –80°C until used. Frozen blocks were sectioned at a thickness of 20 µm using a cryostat (Leica, CM3050S), mounted on Superfrost plus slides (Fisher brand) and washed in PBS supplemented with Triton X-100 (0.1 %, PBST). Unspecific binding sites were blocked with 10% normal goat serum in PBST for 1h at room temperature. Sample were then incubated at 4°C in a solution of 2.5% normal goat serum in PBST using a dilution of 1:500 of the mouse anti acetylated α-tubulin antibody (Sigma, T6793) (Table S3). The primary antibody was then washed with PBST for at least 30 min before incubation with a goat anti-mouse antibody coupled to Alexa-555 (Invitrogen, A-21422).

Finally, samples were stained with a 10 µM DAPI solution (Invitrogen, D3571), dried and mounted in Prolong Gold (Thermo Fisher Scientific). Control sections not exposed to the primary antibody were devoid of immunostaining and were used to set background values on the microscope. Samples were imaged using a Zeiss LSM900 or LSM980 confocal microscope and images analyzed with the ImageJ software (NIH).

### Immunocytochemistry

Cells were cultured in 96-well plates and transfected as described above. Genes were allowed to express for at least 24h and cell fixed with 4% paraformaldehyde in PBS for 1h at room temperature. Samples were blocked with 10% normal goat serum in PBST for 1h at room temperature and then incubated overnight at 4°C with a solution of 2.5% normal goat serum in PBST and a mouse anti-HA-tag or rabbit anti-FLAG-tag primary antibody (1:300). The primary antibody was then washed with PBST followed by incubation of the secondary antibody goat anti-mouse or goat anti-rabbit conjugated to Alexa-555 during 2 h at room temperature. Nuclei were stained with DAPI and the coverslips dried and mounted in Prolong Gold (Thermo Fisher Scientific). Cells were imaged using a Zeiss LSM900 or LSM980 confocal microscope and images analyzed with custom MATLAB software.

### In situ hybridization

Tissues were collected from deeply anesthetized animals, embedded in OCT and immediately frozen in dry ice. Samples were stored at –80°C until used. Tissue was sectioned at a thickness of 20 µm using a cryostat (Leica, CM3050S) and mounted on Superfrost plus slides (Thermo Fisher Scientific). Multiplex fluorescent in situ hybridization was performed using the multiplex v2 RNAscope platform and RNA probe sets were designed by Advanced Cell Diagnostics. The manufacturer’s recommended protocol was followed for probe hybridization and amplification in fresh frozen tissue. Probes were visualized using the fluorescent dyes TSA-FITC, TSA-Cy3 and TSA-Cy5 and nuclei stained using DAPI. Samples were imaged using a Zeiss LSM980 confocal microscope and images analyzed with custom ImageJ (NIH) and MATLAB software.

### Quantitative RT-PCR

Sensory tissue from sucker cups was dissected, flash froze and kept at –80°C until use. Tissue was homogenized with a pestle and RNA extracted with cold TRIzol (Invitrogen) according to the manufacturer protocol. Purity of the isolated RNA was measured in a Nanodrop spectrophotometer. DNase and cDNA synthesis were performed using the enzymes DNase I (EN0521, Thermo) and the high-capacity cDNA reverse transcription kit (Applied Biosystems), respectively. Quantitative real-time PCR was performed using oligo probes (Azenta) (Table S4) and the Kapa SYBR Fast qPCR kit (KAPA Biosystems) on a CFX96 Touch Real-Time PCR detection system (Bio-Rad). Technical duplicates were performed for each qPCR reaction. The relative fold change of gene expression was calculated relative to the housekeeping gene Gapdh. Sequences of all primers are provided in Table S4.

### Scanning electron microscopy

SEM was performed by the Harvard Medical School Electron Microscopy Core Facility and according to standard protocols. Samples were fixed in a solution containing 1.25% formaldehyde, 2.5% glutaraldehyde and 0.03% picric acid in 0.1M sodium cacodylate buffer, pH 7.4 (Electron Microscopy Sciences). Samples were washed with 0.1 M sodium cacodylate buffer and post-fixed with 1% osmium tetroxide in 0.1 M sodium cacodylate buffer for 2 h. Tissues were rinsed in ddH2O and dehydrated through an ethanol series (30, 50, 70, 95 and 100%) for 15 min in each step. Tissues were then dried using an autosamdri-815 critical point dryer and mounted on aluminum stages with silver paint and coated with platinum (10 nm). Mounted tissues were imaged on a Hitachi S-4700 Field Emission Scanning Electron Microscope at an accelerating voltage of 5kV.

### Phylogenetic and Selection analysis

We used the *O. bimaculoides* CRs, Elav and NompC coding sequences as queries for the search of homologue sequences across different species using JackHMMER (Table S5) with an e-value of 10e-10. Sequences were clustered at 100% identity using CD-Hit to collapse duplicates and aligned with MAFFT. Maximum likelihood trees were built using IQ-TREE. For selection analysis, CR and acetylcholine receptor like sequences were mined from cephalopod genomes (Table S6). We generated codon alignments using PAL2NAL, Hyphy and MAFFT. We analyzed positive selection sites using the Mixed Effect Model of Evolution (MEME) from HyPhy with a default threshold of significance of LRT= 4.3. In addition, we analyzed purifying selection pressures with Fixed Effects Likelihood (FEL) from Hyphy.

### Sequencing library preparation

Sensory tissue from octopus sucker cups was mechanically homogenized on ice with a pestle in 1 mL of chilled lysis buffer (10 mM Tris-HCl pH 7.5, 146 mM NaCl, 1 mM CaCl_2_, 21 mM MgCl_2_, 0.03% Tween-20, 0.01% BSA, 1 mM DTT, 10% EZ Lysis Buffer, 0.2–1 U/µL RNase inhibitor). The homogenate was incubated on ice for 5 min, filtered through a 70 µm strainer, and centrifuged at 500 g for 5 min at 4 °C. The pellet was washed twice in nuclei wash buffer (10 mM Tris-HCl pH 7.5, 10 mM NaCl, 3 mM MgCl_2_, 1% BSA, 1 mM DTT, 0.2–1 U/µL RNase inhibitor) and passed through a 40 µm filter to remove debris. Nuclei yield and integrity were assessed using both acridine orange/propidium iodide (AO/PI) and Trypan Blue staining and microscopy. For library construction, nuclei were loaded at the manufacturer-recommended concentration to achieve 10,000 barcoded nuclei per sample using the Chromium Next GEM Single Cell Multiome ATAC + Gene Expression Kit (10x Genomics). Libraries were prepared according to the manufacturer’s protocol and sequenced on an Illumina NovaSeq 6000 S4 flow cell to the recommended depth for Multiome experiments (10x Genomics), ensuring at least 70% sequencing saturation per library.

### Single nuclei RNA sequencing analysis

Illumina sequencing reads were aligned to the *Octopus bimaculoides* genome (GCA_001194135.2) using the 10x Genomics Cell Ranger pipeline with default parameters. We used the R package Seurat and standard data analyses practices. For the hectocotylus sample obtained from the proximal region (base) we filtered nuclei below 100 UMIs, 50 genes and greater than 20% mitochondrial reads. For all other samples we filtered nuclei below 200 UMIs, 100 genes and greater than 10% mitochondrial reads. After filtering, the data were normalized in Seurat using SCTransform function on the 3000 most variable features. We then integrated the data using harmony. To visualize the data post-integration, we calculated the top 40 principal components (PCs) and used these to derive reduced uniform manifold approximation and projection (UMAP) components and UMAP reduction. We tested multiple resolutions for cell clustering and determined the best resolution by examining the expression of known marker genes and looking at specific gene markers for each cluster that were identified with the FindAllMarkers function using the Wilcoxon rank sum test.

### CRT1 purification

The Sleeping Beauty stable cell line for CRT1 expression was generated using a protocol described previously (19). Briefly, cells were grown to 6.4 L at a density of approximately 3.5 - 4.0 × 106 cells per milliliter at 37 °C. Then, 1 µg/mL doxycycline and 1 mM sodium butyrate were added to induce the expression of CRT1 and cells were cultured at 30 °C. Cells were harvested 48 h after induction by centrifugation, resuspended in TBS (20 mM Tris, 150 mM NaCl, pH 7.4) containing 1 mM phenylmethanesulfonyl fluoride (Sigma-Aldrich), then lysed using an Avestin Emulsiflex. Lysed cells were centrifuged for 15 min at 10,000 g and the supernatant was collected and centrifuged again for 2 hr at 186,000 g. The membrane pellet was stored at −80 °C until use.

The membrane pellet was thawed in TBS buffer with 1 mM PMSF and mechanically homogenized using a Dounce homogenizer. The membrane was solubilized for 1 h at 4 °C in TBS buffer with 40 mM n-dodecyl β-D-maltoside (DDM, Anatrace). Solubilized membranes were centrifuged for 40 min at 186,000 g at 4 °C. The supernatant was collected and passed over Strep-Tactin affinity resin (2 ml) at 0.8 ml/min. The resin was washed with TBS buffer containing 1 mM DDM, then eluted with elution buffer (20 mM Tris, 150 mM NaCl, pH 8.0) supplemented with 5 mM D-desthiobiotin (Sigma-Aldrich) and 1 mM DDM. The CRT1 protein was concentrated to approximately 500 µL for nanodisc reconstitution.

### Nanodisc reconstitution

The plasmid for saposin A expression was obtained from Salipro Biotech AB (29). Reconstitution was carried out at a molar ratio of protein, saposin, and soy polar lipids of 1:50:250, with the addition of 0.2 mM progesterone. The mixture was assayed by tryptophan fluorescence SEC and concentrated, then centrifuged for 20 min at 40,000 rpm to remove aggregates before size-exclusion chromatography (SEC). The resulting supernatant was separated by SEC using a Superose 6 Increase 10/300 GL column (Cytiva) equilibrated in TBS buffer at pH 8.0 containing 200 µM progesterone in 1% DMSO. SEC fractions corresponding to pentameric CRT1-progesterone were pooled and concentrated and the quality of final sample was assessed by tryptophan fluorescence SEC before freezing grids.

### Cryo-EM sample preparation

A total of 3 µL of concentrated sample was applied to copper R1.2/1.3 300 mesh holey carbon grids (Quantifoil) that were glow-discharged at 30 mA for 80 s. The grids were immediately blotted for 2.5 s under 100% humidity at 4 °C and then plunge-frozen into liquid ethane cooled by liquid nitrogen using a Vitrobot Mark IV (Thermo Fisher Scientific).

### Cryo-EM data collection and processing

After screening grids in the UCSD Cryo-EM facility, grids were sent to the Pacific Northwest Center for Cryo-EM for large dataset collection. In total, 7,517 dose-fractioned images were collected on a 300 kV Titan Krios 1 (Thermo Fisher Scientific) at the PNCC using a Falcon 4i direct electron detector with Selectris X energy filter. The total exposure was 50 e-/Å2 and the defocus range was set to −0.8 µm to −2.0 µm with a magnification of 130,000× and 0.935 Å per pixel. The data processing was done using cryoSPARC v.4.6 and v.4.7 (30). Dose-fractionated images were gain-normalized, motion corrected and CTF estimated in a CryoSPARC live session with default settings. Particles were picked using the template picker with a diameter of 200 Å and extracted with a 320-pixel box size at bin 2. After several rounds of two-dimensional (2D) classification, ab initio and heterogeneous refinement were used to filter the junk particles. Selected particles were re-extracted at full size and subjected to non-uniform (NU) refinement (31) to generate the 3D volume with C5 symmetry. The final map for CRT1-progesterone is at a resolution of 2.77 Å with 241,406 particles (Table S7).

### Model building, refinement and validation

PDB 8EIS (18) was used as the starting model. The starting model was aligned and fitted into the EM density map in UCSF ChimeraX (v.1.9) (32), followed by several rounds of manual building in Coot (v.0.9.8.93) and global real space refinement in Phenix (v.1.20.1) (33) using secondary structure restraints and Ramachandran restraints. Based on the ligand density, four different poses of progesterone were modeled, and molecular dynamics (MD) simulations were subsequently performed for each model (see details below). The most favorable pose was identified as the one exhibiting the lowest ligand root-mean-square deviations (RMSD) value. Model geometry and clash scores were checked by Molprobity (34). The ligand restraint files were generated using the Grade Web Server (https://grade.globalphasing.org/) with default settings.

Figures of protein structures and density maps were generated using UCSF ChimeraX (v.1.9) and PyMOL (v.2.5.5, Schrodinger, LLC). The pore radius profiles were calculated by HOLE v.2 (35), and the hydrophobicity of the permeation pathway was generated by CHAP (36). The 2D ligand interaction diagrams show residues within 5 Å of the ligand and was generated using LigPlot+ v.2.2.9 (37). The interaction area between ligands and CRT1, as well as the solvation energy of related residues, were calculated by PDBePISA (38).

### Molecular dynamics simulations

Molecular dynamics simulations of progesterone bound to CRT1 were performed using GROMACS-2024.2 (39) utilizing CHARMM36m (40) force field. The force field parameters for progesterone were generated using the CHARMM General Force Field (CGenFF) (41). Atomics coordinates of CRT1 complex with 5 progesterone molecules were used as the starting model. The model was inserted into 1-palmitoyl-2-oleoylphosphatidylcholine (POPC) lipid bilayers, solvated with water (TIP3P63) and supplemented with 0.3 M NaCl using CHARMM-GUI Membrane Builder (42). Simulations were performed at 303 K and 1 bar using the velocity-rescaling thermostat (43) and stochastic cell rescaling barostat (44). The LINCS algorithm was used to constrain the length of all bonds involving hydrogens (45), and the particle mesh Ewald method (46) was used to calculate long-range electrostatic interactions. The systems were energy minimized and then equilibrated, with the position restraints on the protein and progesterone gradually released. Because the M4 transmembrane helix was unresolved, we retained restraints on the transmembrane domain during production simulations to prevent artefacts from the missing M4 helix. Six replicates 100 ns each were simulated as final production runs. MD simulation trajectories analysis was performed using MDAnalysis (47). RMSD of progesterone atomic coordinates were calculated in MDAnalysis (47) and plotted as violin plots using Matplotlib (48).

### Quantification and statistical analysis

All data were analyzed and plotted with custom MATLAB software (MathWorks) or RStudio (Posit Software, PBC). Data are represented as mean ± SEM and n values represent independent experiments. Data were significant if p-values were below a 0.05 threshold using a non-parametric one-way Anova with a Kruskal-Wallis test. All i*n vivo* and *ex vivo* animal behavior trials were randomized.

**Figure S1.**
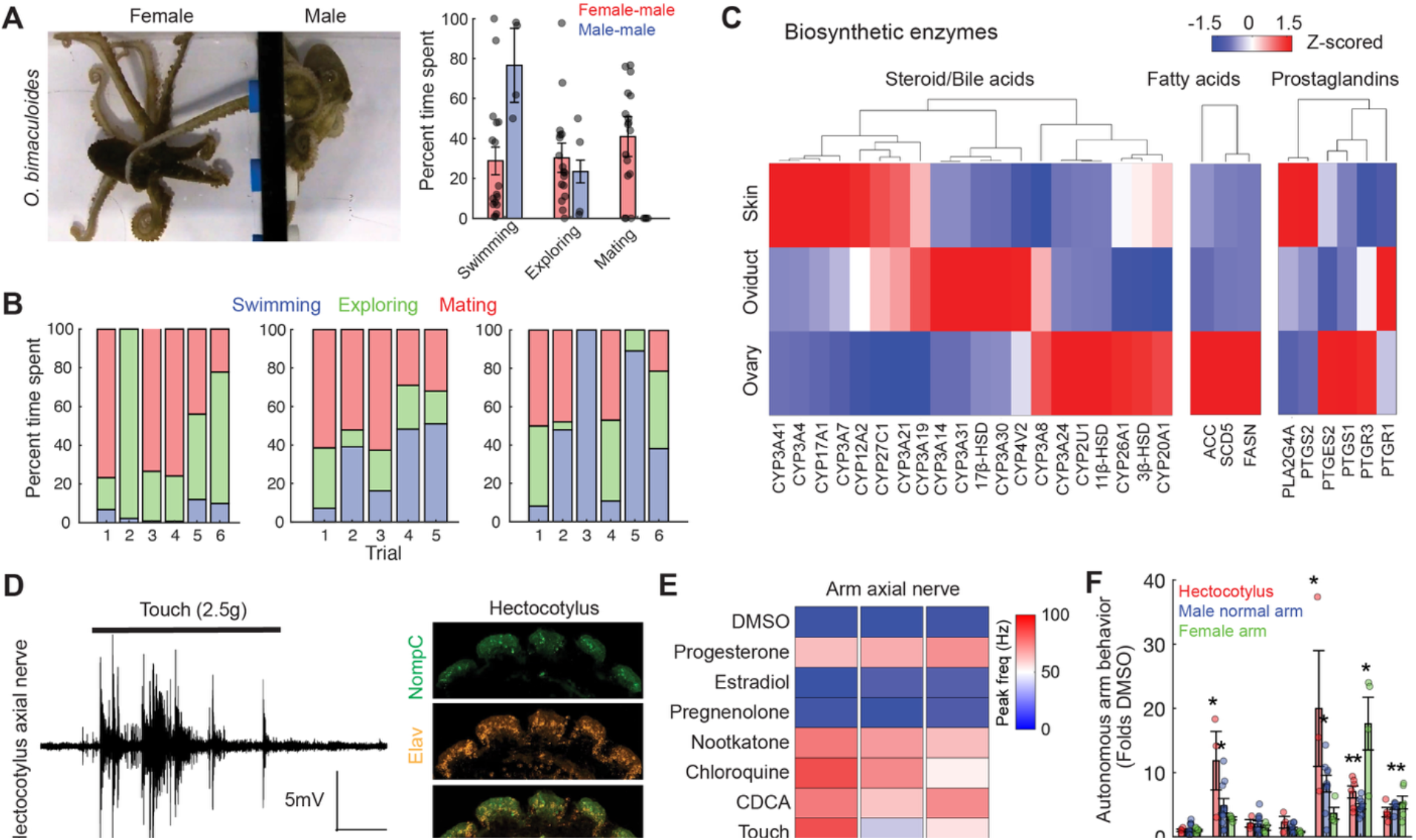
Chemosensory mating in octopus. **(A)** Left, overhead view of the mating behavior arena. Octopuses interacted through a black, opaque acrylic separation with three holes 8 mm in diameter at the bottom of a 60×20×25 cm tank. At the start of each trial, octopuses swam until they reached through the wall to initiate the exploration phase. Mating was defined as the period when the male introduces its hectocotylus into the other octopus’s mantle. Right, summary of percent time spent mating for female-male (n= 3) and male-male (n= 3) couples. No male-male mating occurred (n= 4–14, p= 0.003, one-way ANOVA). **(B)** Mating occurred in all tested pairs and in most trials. Trial-by-trial examples for three wild caught female-male *O. bimaculoides* mating pairs. Trials were about one hour long and were carried out in consecutive days or every other day. **(C)** RNA-seq expression profile of biosynthetic enzymes expressed in the mantle skin, oviductal gland, ovary and brain. Most abundant synthetic pathways found correspond to the production of fatty acids, prostaglandins and steroid/bile acids. Data was z-scored across 3 samples. **(D)** Left, touch elicited axial nerve activity from the hectocotylus, similar to normal arms (female arm: 62.5 ± 14Hz, n= 4; male normal arm: 24 ± 6Hz, n= 3; hectocotylus: 37.5 ± 15Hz, n= 4). Right, the hectocotylus tip expresses the mechanoreceptor NompC (No mechanoreceptor potential C, green), as observed by in situ hybridization. The neuronal marker Elav is shown in red (n= 3). **(E)** Steroids evoked neural responses in hectocotylus and normal arms: hectocotylus 55 ± 14 Hz compared to baseline, n = 3; male arms 47.7 ± 3 Hz, n = 7; female arms 45.3 ± 9 Hz, n= 6. **(F)** Summary data for amputated arm behavioral responses for steroids, bile acids and bitter molecules in *Octopus bimaculoides* from Fig. 1C. Hectocotylus, n= 3–6; normal male arm, n= 5–10; female arms, n= 4–6. Statistical comparisons with DMSO were calculated using one-way ANOVA with Kruskal-Wallis test. p< 0.005.

**Figure S2.**
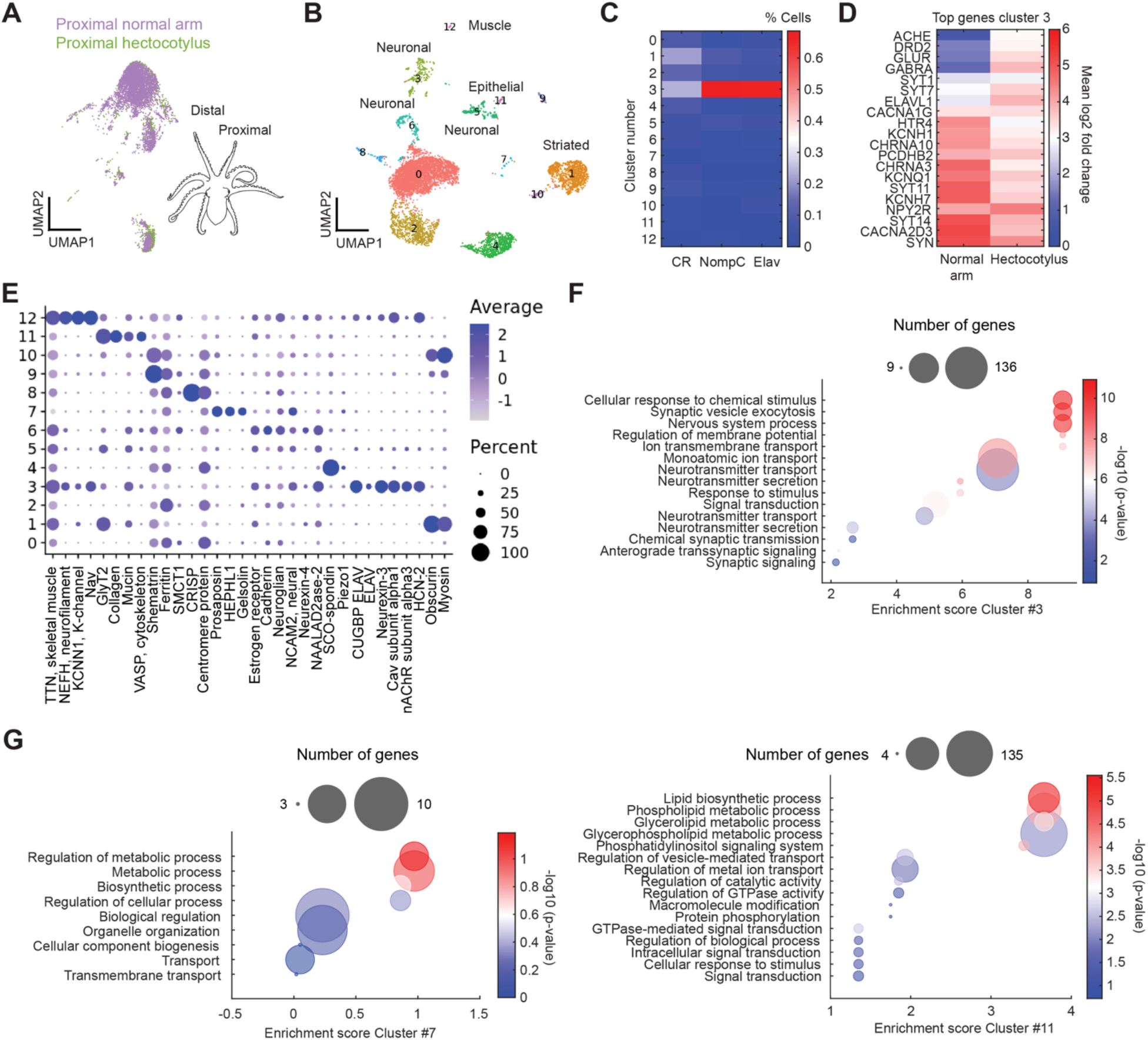
snRNA-seq analysis of the octopus hectocotylus. **(A)** UMAP reduction of snRNA-seq data from sucker sensory epithelium in the arm base (proximal) of the normal arm and hectocotylus shows an overlay of the cell population for the two samples indicating similar transcriptional programs (1502 nuclei for hectocotylus base versus 4875 cells for normal arm base). **(B)** UMAP analysis of clustered cell populations from the normal arm tip (distal). A total of 2702 single nuclei were sequenced and shown colored by clusters. Clusters ranged from 22 to 1189 nuclei and identity was inferred by expression of enriched marker transcripts. **(C)** Percent of cells expressing CRs, NompC or the neuronal marker Elav over the subpopulation of cells where these transcripts were detected and across all clusters. Cluster 3 showed highest number of Elav (68.5%), NompC (66.2%) and CR (23.9%) expressing cells across clusters. **(D)** Comparison between normal arm and hectocotylus for the highest expressed transcripts in Cluster 3 shown as mean log2 fold change, where 0 represents no change and 1 a two-fold higher expression. The dopamine receptor D2, glutamate receptor and GABA receptor, among others, have higher expression levels in the hectocotylus compared to normal arms. **(E)** Top enriched genes across all clusters. Clusters 3, 6 and 7 express genes associated to neuronal function. **(F)** Gene ontology analysis (GO) for cluster 3 shows “cellular response to chemical stimulus” as the most enriched term. Bubble plot represents the number of transcripts per category (bubble size = number of genes) versus the enrichment score and significance (p-value) obtained using the DAVID gene annotation tool (NCBI). **(G)** Gene-ontology of non-neuronal Cluster 7 and cluster 11 which have higher numbers of nuclei in the hectocotylus compared to normal arm (related to Fig. 2F). Cluster 7: hectocotylus tip 3.6% vs. normal arm tip 0.5% from all nuclei. Cluster 11: hectocotylus tip 1.6% vs. normal arm tip 0.4% from all nuclei.

**Figure S3.**
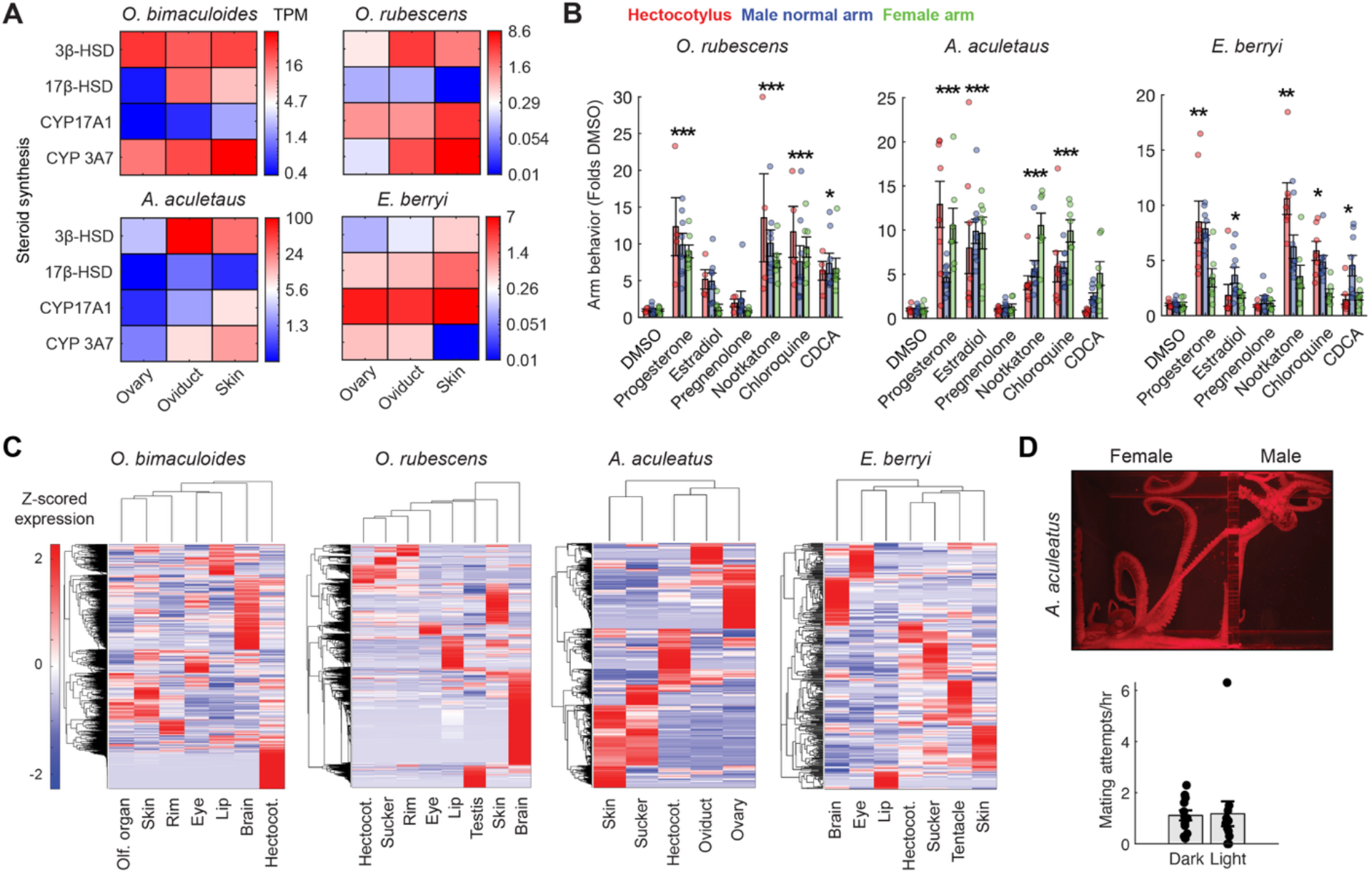
Comparative behavior and transcriptomics in octopus and squid species. **(A)** Steroid biosynthetic enzymes were expressed in the ovary, oviduct and skin of the octopuses *O. bimaculoides, O. rubescens, A. aculeatus* and squid *E. berryi*. The color scale shows raw transcripts per million (TPM) for each sample. **(B)** Summary data for normal arm (blue), female arm (green) and hectocotylus (red) behavioral responses for steroids, bile acids and bitter molecules in *O. rubescens* (n= 4–8), *A. aculeatus* (n= 8–9) and *E. berryi* (n= 7–9) from Fig. 3C. Statistical comparisons with DMSO were calculated using one-way ANOVA with Kruskal-Wallis test. p< 0.005. **(C)** Tissue-based transcriptomics across sensory tissues for octopus and squid shows distinct gene expression programs for hectocotylus and normal arms in all species. **(D)** Similar to *O. bimaculoides*, the divergent species *A. aculeatus* mates in the absence of visual cues (n= 7 mating pairs, dark 0.68 ± 0.14 vs. light 0.66 ± 0.27 mating attempts per hour), suggesting chemosensory mating is widespread across the octopus phylogeny.

**Figure S4.**
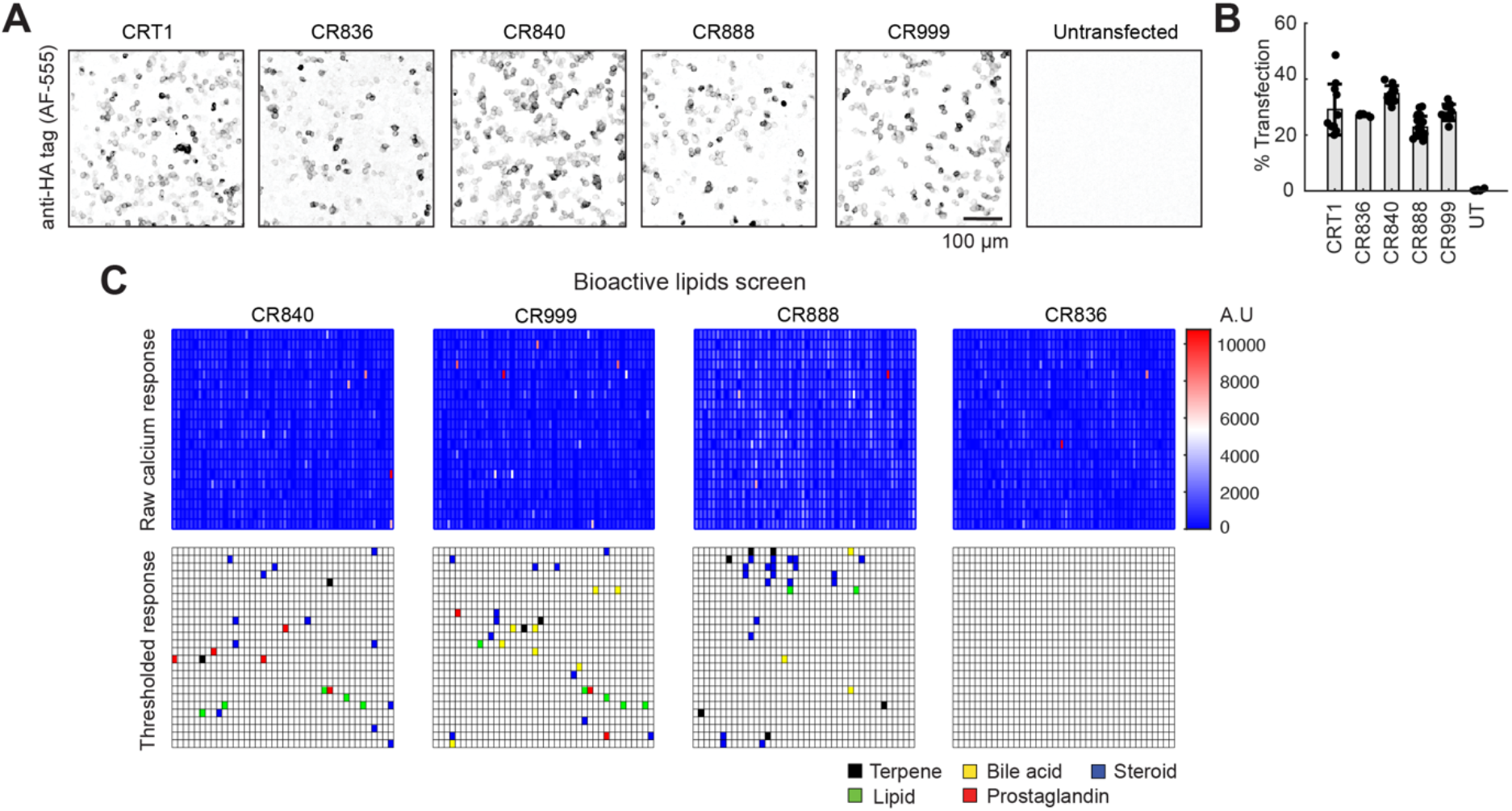
Bioactive lipid screen in octopus CRs. **(A)** Octopus CRs were expressed in HEK293 cells together with the calcium sensor GCaMP6s to monitor receptor activity. Representative images of hemagglutinin (HA)-tagged CRs labeled using an anti HA-tag antibody to validate protein expression. Immunostaining was absent in untransfected cells. **(B)** The transfection rate was estimated as the percentage of immunostained cells over the total number of DAPI cells. Each dot represents mean transfection percentage in a single image. **(C)** Thresholded responses in HEK cell expressing a given CR (top label) in response to molecules in the bioactive lipid panel (see Table S2). Each cell represents the maximal fluorescence after the addition of a single compound at a 10 µM final concentration. Each screen plate contained 8 control wells (vehicle) used for comparison with test wells. Responses greater than the mean vehicle response plus 8 times the standard deviation across vehicle wells were considered significant and assigned to a particular category based on molecular descriptors.

**Figure S5.**
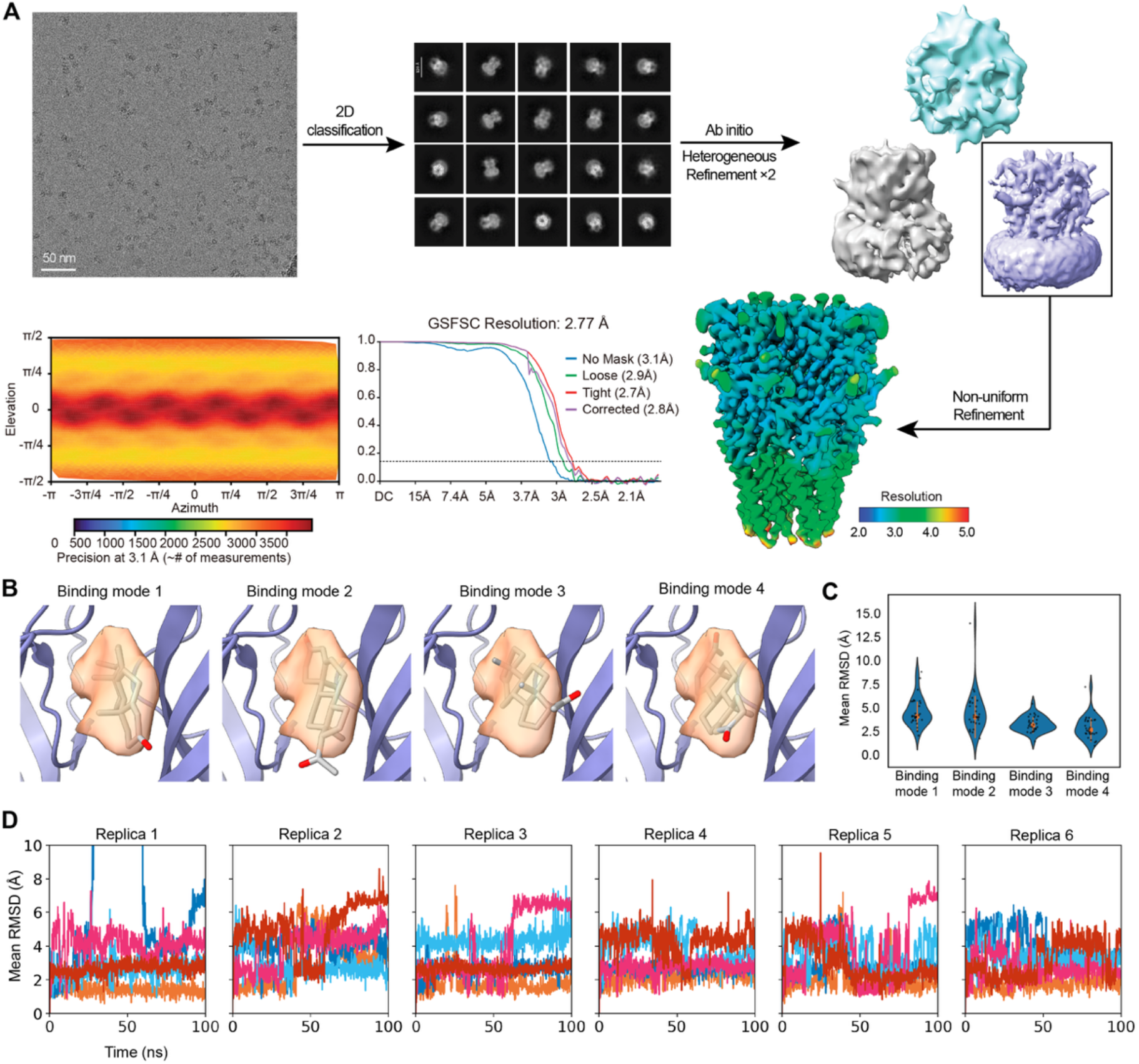
Cryo-EM data processing and model building. **(A)** Representative cryo-electron micrograph of CRT1 plus progesterone sample. Particles were subjected to two-dimensional classification followed by ab-initio and heterogeneous refinement to generate an initial model. After nonuniform refinement, a ∼2.8 Å resolution reconstruction was obtained with C5 symmetry. **(B)** Atomic model of CRT1 ligand binding pocket showing progesterone in 4 binding modes based on cryo-EM density. Mode 1: methyl ketone inside, two methyl groups left. Mode 2: methyl ketone outside, two methyl groups right. Mode 3: methyl ketone outside, two methyl groups right. Mode 4: methyl ketone inside, two methyl groups right. **(C)** Progesterone binding mode stability in MD, quantified by ligand root-mean-square deviations (RMSD) (Å). Each point represents the mean RMSD over a 100 ns simulation for a single progesterone molecule. Violin plots depict the distribution of mean RMSD per binding mode. The orange marker indicates group median and error bars denote ±1 SD. Simulations of CRT1 protein with 5 progesterone molecules bound were run in 6 independent replicates (n= 30 per binding mode). Mode 4 exhibits the lowest RMSD, indicating it represents the most energetically favorable pose. **(D)** Progesterone binding mode 4 stability over time. Each trace shows RMSD (Å) for a single progesterone molecule (n=5 per replicate).

**Figure S6.**
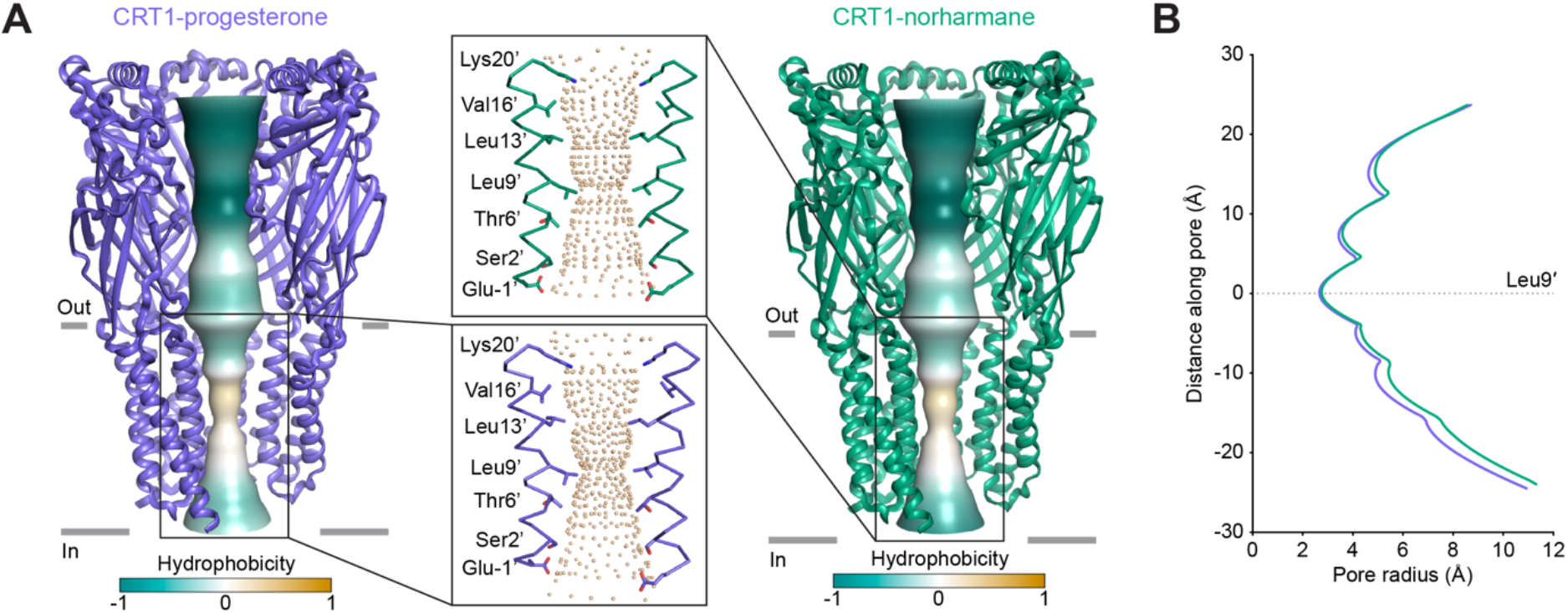
Receptor ion permeation comparison for mating and prey molecules. **(A)** The ion permeation pathway of the CRT1-progesterone (slate blue) and CRT1-norharmane complexes (PDB code: 9E6B, green) colored by hydrophobicity. Inset: Two M2 helices with pore-lining residues are shown as sticks; spheres indicate pore shape. **(B)** Comparison of pore diameters between CRT1-progesterone and CRT1-norharmane, indicating that the CRT1 pore adopts a similarly open conformation.

**Figure S7.**
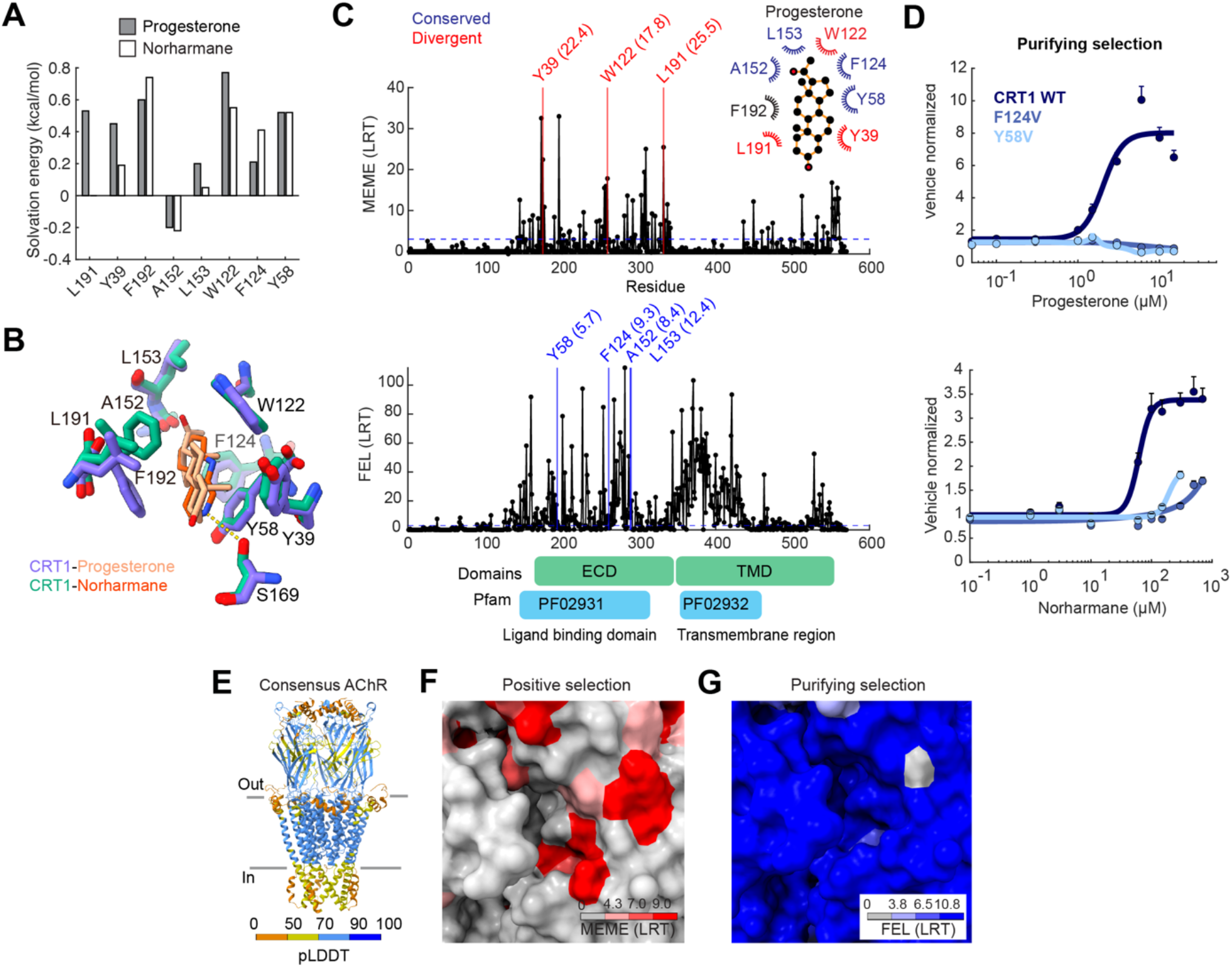
Selection analysis of CRT1 binding pocket. **(A)** Solvation energy of residues L191, Y39 and L153 with progesterone (black) is higher compared to norharmane (white) indicating a more thermodynamically favorable interaction. **(B)** Atomic model of CRT1 ligand binding pocket bound to progesterone (slate blue) or the prey-derived molecule norharmane (green). **(C)** Likelihood ratio test (LRT) for diversifying selection (MEME, top) indicates that ligand interaction residues L191, Y39 and W122 in CRT1 binding pocket are under positive selection, while A152, L153, F124 and Y58 are under purifying selection (FEL, bottom). Threshold for significance was LRT= 4.3 (p<0.05). **(D)** Mutating the conserved residues F124 and Y58 impairs both progesterone and norharmane responses compared to the wild type CRT1. Dose response curves for CRT1 point mutants F124V and Y58V compared to the wildtype receptor for progesterone (top) and norharmane (bottom). **(E)** Predicted AlphaFold3 (23) structure of a consensus octopus nicotinic acetylcholine receptor (AChR). A consensus protein sequence was obtained from 299 AChR sequences across 30 cephalopod species including octopus, squid and cuttlefish species. **(F)** Residues in the binding pocket of a consensus AChR colored by the likelihood ratio test (LRT) value for the positive selection pressure test MEME. Threshold for significance, LRT= 4.3 (p<0.05), LRT= 7 (p<0.01), LRT= 9 (p<0.001). **(G)** Residues in the AChR binding pocket colored by the likelihood ratio test (LRT) value for the purifying selection pressure test FEL. Threshold for significance, LRT= 3.8 (p<0.05), LRT= 6.5 (p<0.01), LRT= 10.8 (p<0.001).

**Figure S8.**
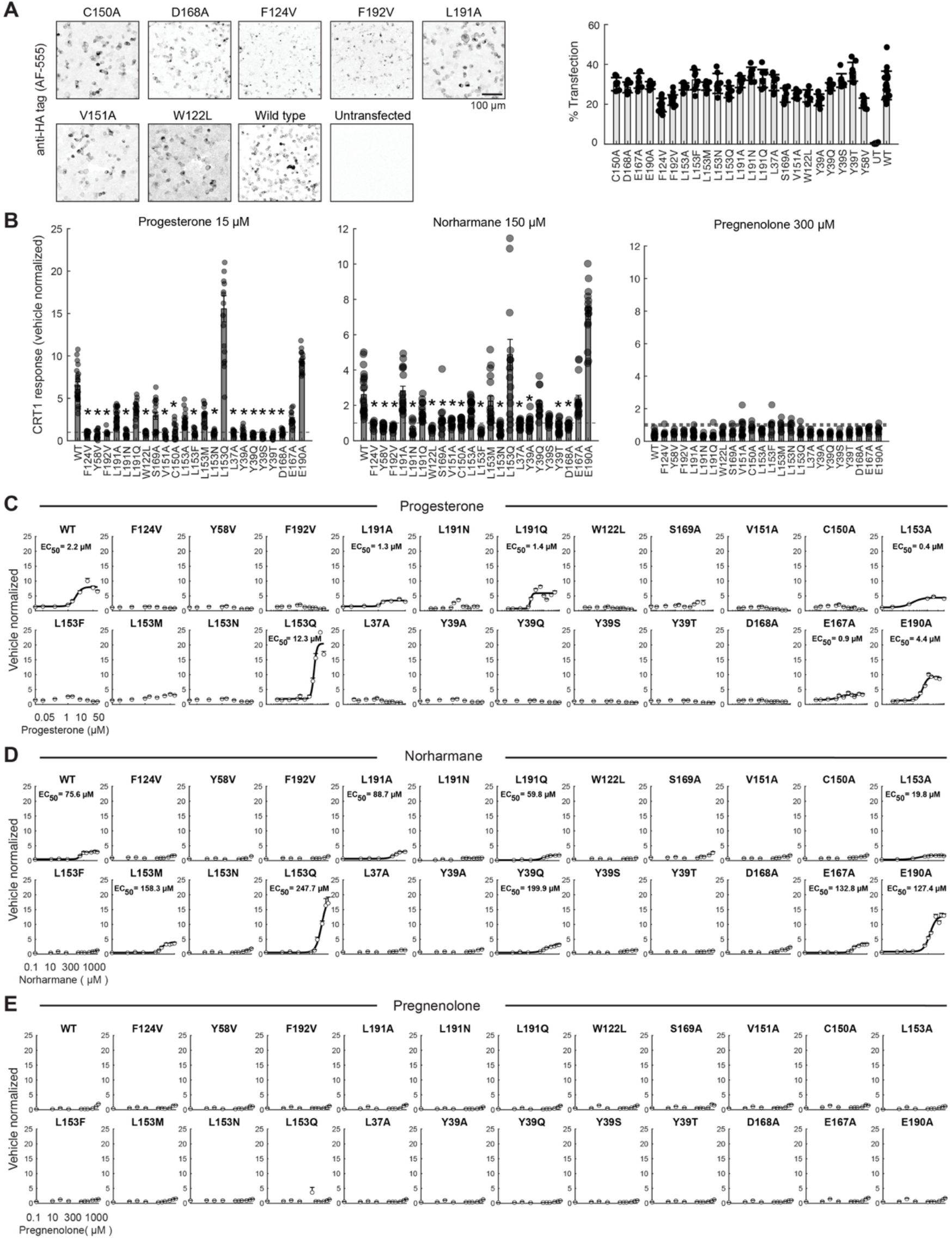
CRT1 mutagenesis. **(A)** Wild type (WT) CRT1 and mutants were tagged with HA their expression in HEK293 cells was verified using immunocytochemistry. WT (n= 19), C150A (n= 8), D168A (n= 8), E167A (n= 8), E190A (n= 8), F124V (n= 12), F192V (n= 16), L37A (n= 8), L153A (n= 8), L153F (n= 8), L153M (n= 8), L153N (n= 8), L153Q (n= 8), L191A (n= 8), L191N (n= 8), L191Q (n= 8), S169A (n= 8), V151A (n= 8), W122L (n= 8), Y39A (n= 8), Y39Q (n= 8), Y39S (n= 8), Y39T (n= 8), Y58S (n= 8), Y58V (n= 8), untransfected (UT, n= 8). p< 0.001, one-way ANOVA with Kruskal-Wallis test compared with UT cells. **(B)** Calcium responses to progesterone (left), norharmane (center) or pregnenolone (right) of WT CRT1 was affected by most point mutations of residues involved in the coordination of the listed ligand. WT (n= 8–32), F124V (n= 8–16), Y58V (n= 8–16), F192V (n= 8–16), L191A (n= 15–24), L191N (n= 8–16), L191Q (n= 8–16), W122L (n= 8–16), S169A (n= 8–16), V151A (n= 15–16), C150A (n= 8– 16), L153A (n= 8–24), L153F (n= 8–16), L153M (n= 16), L153N (n= 15–16), L153Q (n= 16), L37A (n= 8–16), Y39A (n= 8–24), Y39Q (n= 8–16), Y39S (n= 8–16), Y39T (n= 8–16), D168A (n= 8–16), E167A (n= 8–16), E190A (n= 8–16). p< 0.001, one-way ANOVA with Kruskal-Wallis test. **(C)** Concentration dependence of progesterone responses in WT CRT1 versus point mutants. n. r= no response. WT (EC50= 2.2 ± 0.2 µM, n= 24), F124V (n. r, n= 16), Y58V (n. r, n= 16), F192V (n. r, n= 8), L191A (EC50= 1.3 ± 0.1 *µ*M, n= 24), L191N (n. r, n= 16), L191Q (EC50= 1.4 ± 0.1 *µ*M, n= 16), W122L (n. r, n= 16), S169A (n. r, n= 16), V151A (n. r, n= 16), C150A (n. r, n= 16), L153A (EC50= 0.4 ± 0.0 *µ*M, n= 24), L153F (n. r, n= 16), L153M (n. r, n= 16), L153N (n. r, n= 15), L153Q (EC50= 12.3 ± 0.5 *µ*M, n= 16), L37A (n. r, n= 16), Y39A (n. r, n= 24), Y39Q (n. r, n= 16), Y39S (n. r, n= 16), Y39T (n. r, n= 16), D168A (n. r, n= 16), E167A (EC50= 0.9 ± 0.8 *µ*M, n=16), E190A (EC50= 4.4 ± 0.3 *µ*M, n= 16). **(D)** Concentration dependence of norharmane responses in WT CRT1 versus point mutants. n. r= no response. WT (EC50= 75.6 ± 6.6 *µ*M, n= 32), F124V (n. r, n= 8), Y58V (n. r, n= 16), F192V (n. r, n= 16), L191A (EC50= 88.7 ± 5.2 *µ*M, n= 16), L191N (n. r, n= 8), L191Q (EC50= 59.8 ± 3.1 *µ*M, n= 16), W122L (n. r, n= 16), S169A (n. r, n= 16), V151A (n. r, n= 16), C150A (n. r, n= 16), L153A (EC50= 19.8 ± 2.0 *µ*M, n= 24), L153F (n. r, n= 8), L153M (EC50= 158.3 ± 7.6 *µ*M, n= 16), L153N (n. r, n= 16), L153Q (EC50= 247.7 ± 19.4 *µ*M, n= 16), L37A (n. r, n= 16), Y39A (n. r, n= 24), Y39Q (EC50= 200 ± 26.3 *µ*M, n= 12), Y39S (n. r, n= 12), Y39T (n. r, n= 16), D168A (n. r, n= 16), E167A (EC50= 132.8 ±5.7 *µ*M, n=16), E190A (EC50= 127.4 ± 4.7 *µ*M, n= 16). **(E)** Neither WT CRT1 nor mutants responded to pregnenolone. WT (n= 23), F124V (n= 7), Y58V (n= 7), F192V (n= 7), L191A (n= 15), L191N (n= 8), L191Q (n= 8), W122L (n= 7), S169A (n= 7), V151A (n= 15), C150A (n= 8), L153A (n= 16), L153F (n= 8), L153M (n= 8), L153N (n= 8), L153Q (n= 8), L37A (n= 8), Y39A (n= 8), Y39Q (n=8), Y39S (n= 8), Y39T (n= 8), D168A (n= 8), E167A (n=8), E190A (n= 8).

**Table S1.** List with chemicals used for *in vitro* screening of CRs.

| Chemical | Source | Catalog |
| --- | --- | --- |
| Dimethyl sulfoxide | Millipore Sigma | Cat# 276855 |
| Agarose | VWR | Cat# N605 |
| Progesterone | Millipore Sigma | Cat# P8783 |
| 17β-estradiol | Millipore Sigma | Cat# 3301 |
| Pregnenolone |  |  |
| Nootkatone | Sigma-Aldrich | Cat# W316620 |
| Chloroquine | Sigma-Aldrich | Cat# C6628 |
| Chenodeoxycholic acid | Millipore Sigma | Cat# C9377 |
| Androsterone | Cayman | Cat# 15872 |
| Estrone | Cayman | Cat# 10006485 |
| Dehydroepiandrosterone | Cayman | Cat# 15728 |
| 17α, 20 β-Dihydroxy-4-pregnen-3-one | Cayman | Cat# 16146 |
| Medroxyprogesterone | Cayman | Cat# 24908 |
| Prostaglandin F2α diethyl amide | Cayman | Cat# 16023 |
| Prostaglandin F2β | Cayman | Cat# 16410 |
| Prostaglandin D2 dopamine | Cayman | Cat# 9002504 |
| 16,16-dimethyl Prostaglandin A1 | Cayman | Cat# 10080 |
| 15-keto Prostaglandin A1 | Cayman | Cat# 9000142 |
| 15(S)-15-methyl Prostaglandin F2α methyl ester | Cayman | Cat# 16744 |
| Allocholic Acid | Cayman | Cat# 30415 |
| Tauro-Obeticholic Acid | Cayman | Cat# 38878 |
| Dehydrolithocholic Acid | Cayman | Cat# 29544 |
| Murideoxycholic Acid | Cayman | Cat# 20290 |
| Ursodeoxycholic Acid | Cayman | Cat# 31016 |
| Bioactive lipid I Screening Library | Cayman | Cat# 10506 |

**Table S2.**
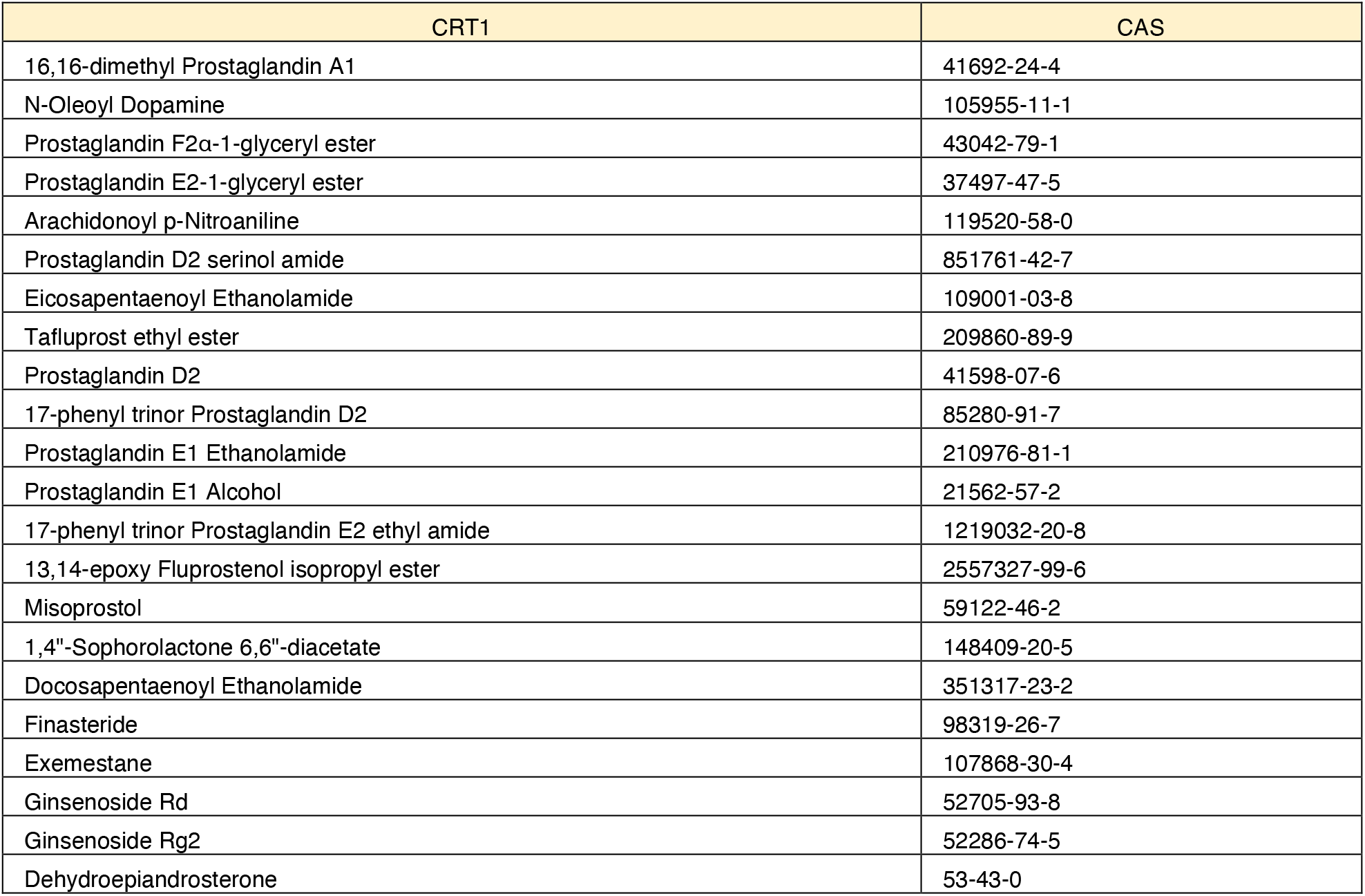
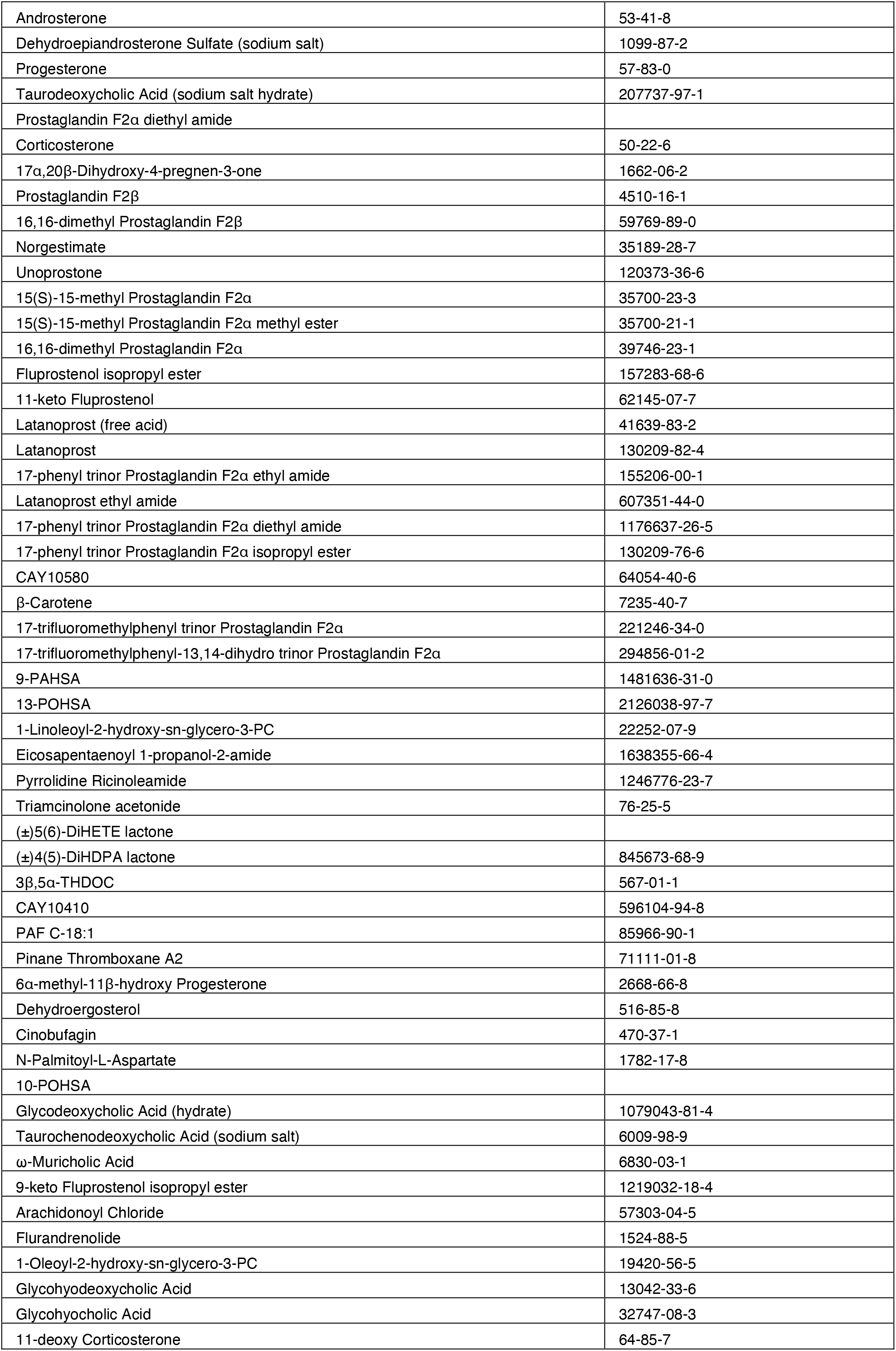
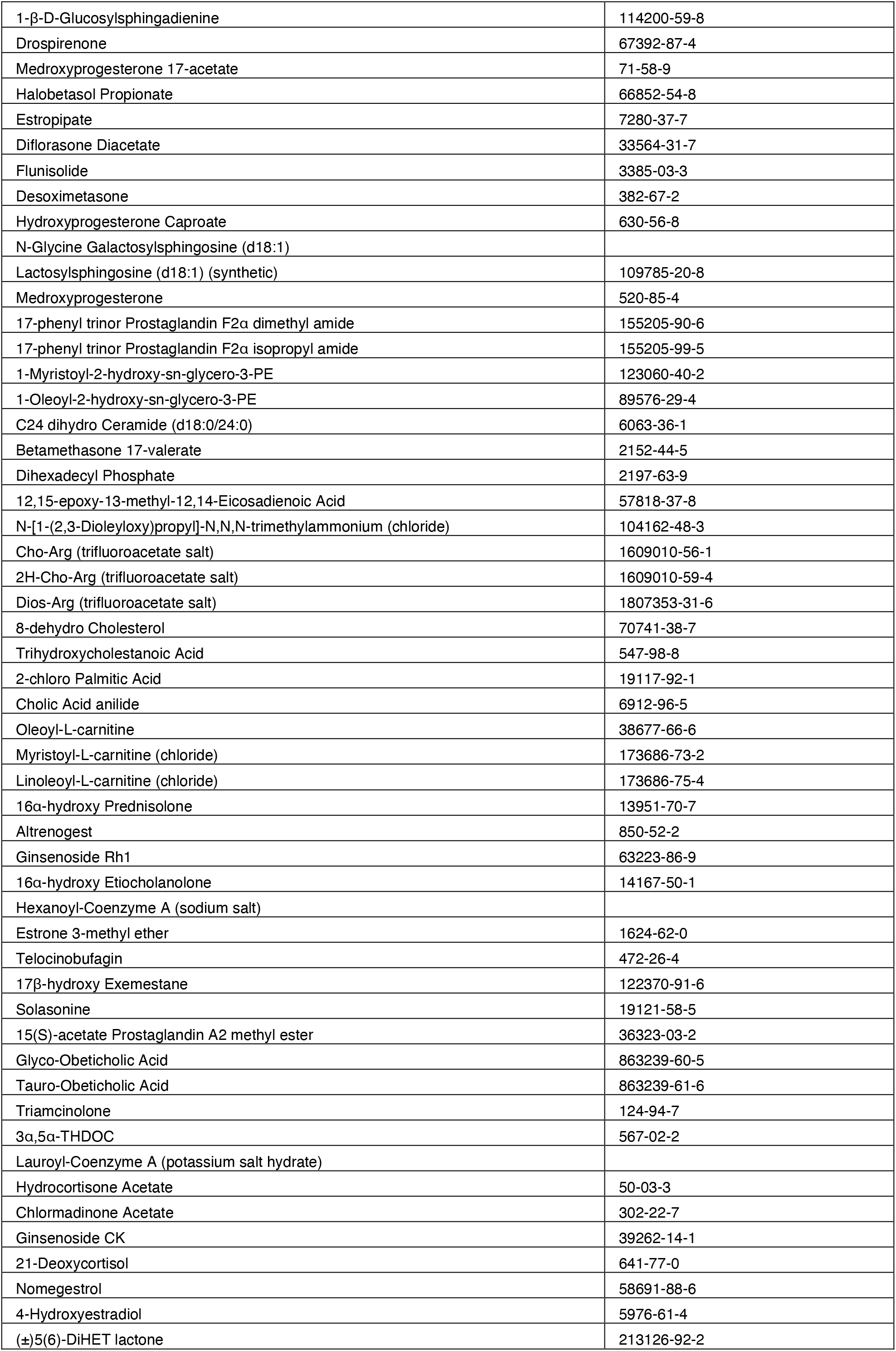
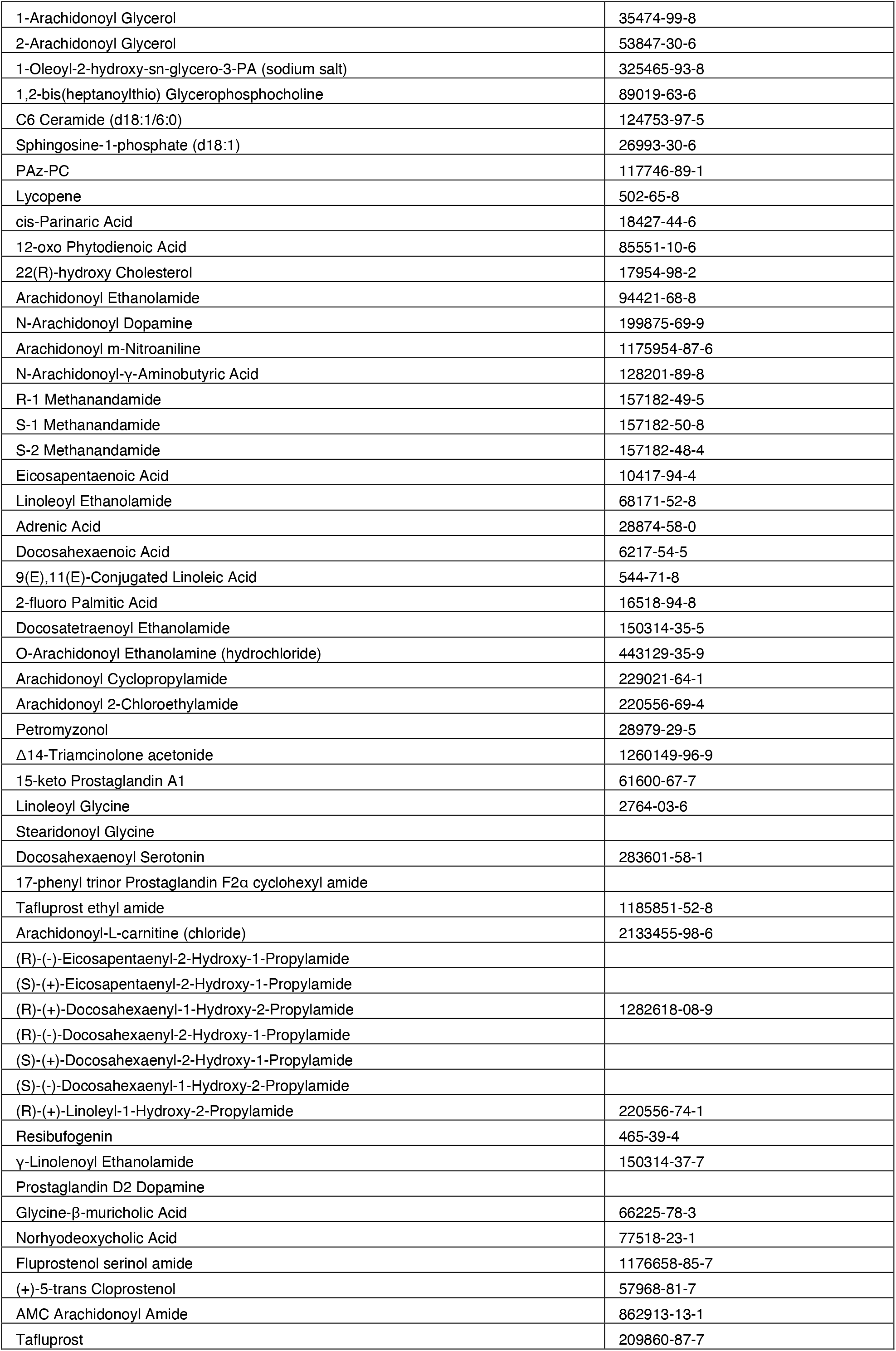
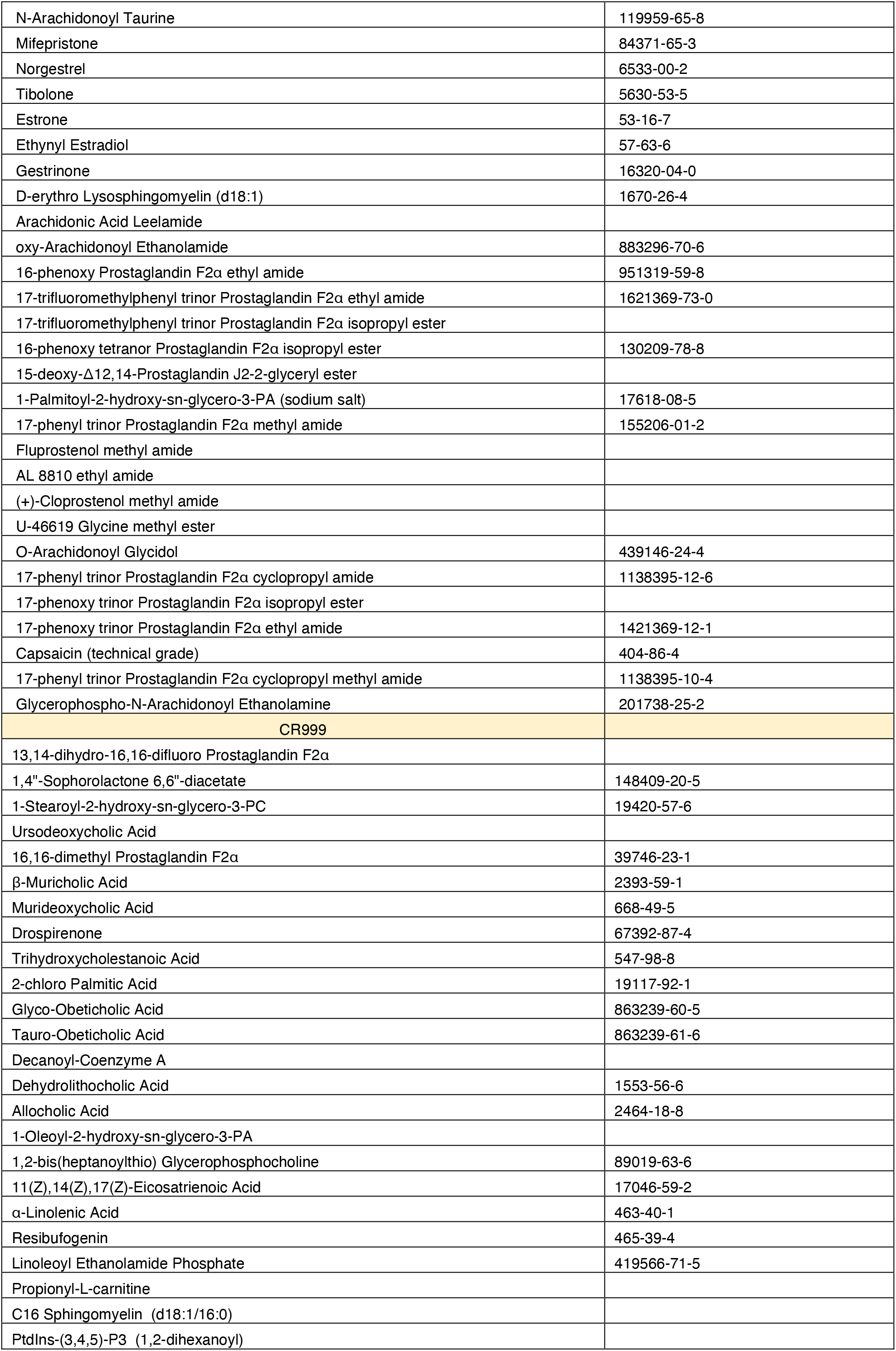
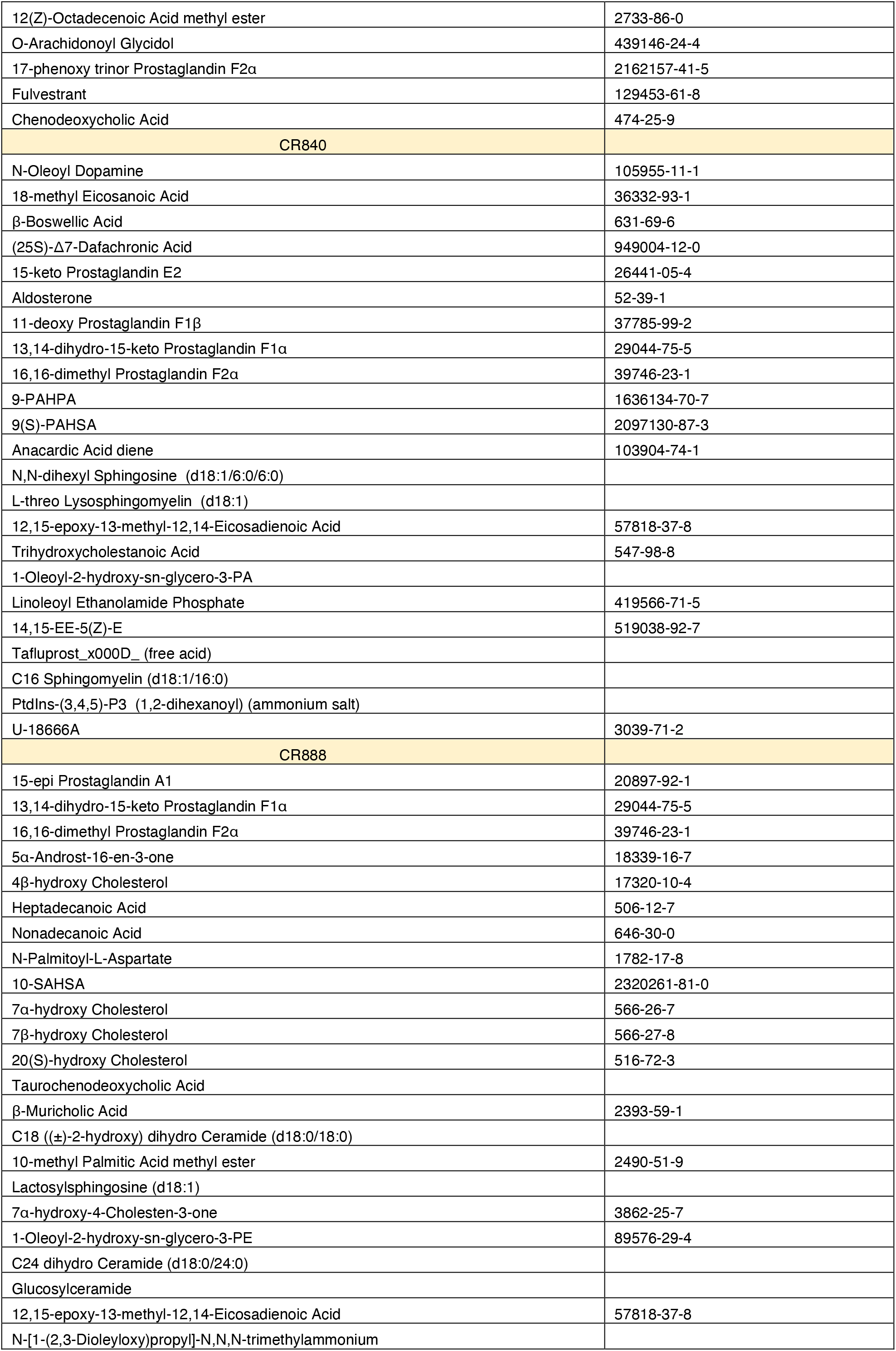
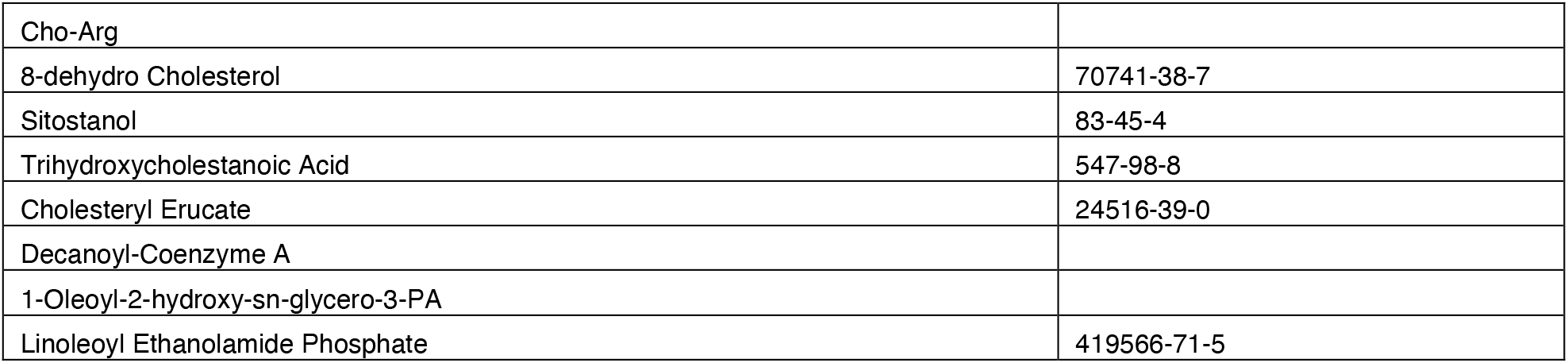
List of molecules that activated each receptor in the active biolipid panel screen and their chemical abstracts service (CAS) registry number, if available.

**Table S3.** List with all antibodies used for immuno staining and references for plasmids used for *in vitro* transfection of wild type CRs and mutagenesis.

| REAGENT | Source | Catalog |
| --- | --- | --- |
| **Antibodies** |  |  |
| Monoclonal Mouse anti-HA | ThermoFisher | 26183 |
| Monoclonal Mouse anti-Tubulin, acetylated | Millipore Sigma | T6793 |
| Goat anti-Mouse IgG, Alexa Fluor 555 | ThermoFisher | A-21422 |
| **Recombinant DNA** |  |  |
| pUNIV CRT1-HA | Van giesen et al., 2020 | N/A |
| pUNIV CR836-HA | This paper | N/A |
| pUNIV CR840-HA | Van giesen et al., 2020 | N/A |
| pUNIV CR888-HA | This paper | N/A |
| pUNIV CR999-HA | This paper | N/A |
| pUNIV CRT1-HA C150A | This paper | N/A |
| pUNIV CRT1-HA F124N | Allard et al., 2023 | N/A |
| pUNIV CRT1-HA F124V | Allard et al., 2023 | N/A |
| pUNIV CRT1-HA F192V | This paper | N/A |
| pUNIV CRT1-HA L153A | This paper | N/A |
| pUNIV CRT1-HA L191A | This paper | N/A |
| pUNIV CRT1-HA L37A | This paper | N/A |
| pUNIV CRT1-HA S169A | This paper | N/A |
| pUNIV CRT1-HA V151A | This paper | N/A |
| pUNIV CRT1-HA W122L | Allard et al., 2023 | N/A |
| pUNIV CRT1-HA Y39A | This paper | N/A |
| pUNIV CRT1-HA Y58S | Allard et al., 2023 | N/A |
| pUNIV CRT1-HA Y58V | Allard et al., 2023 | N/A |

**Table S4.**
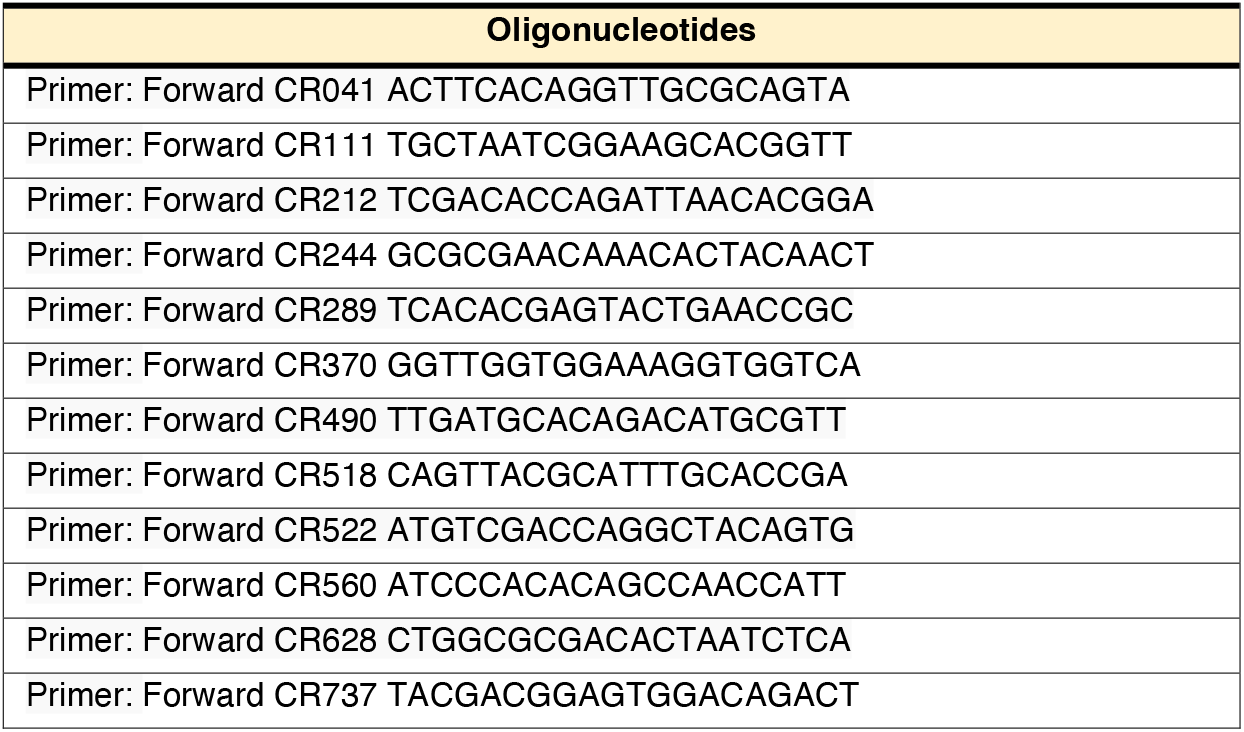
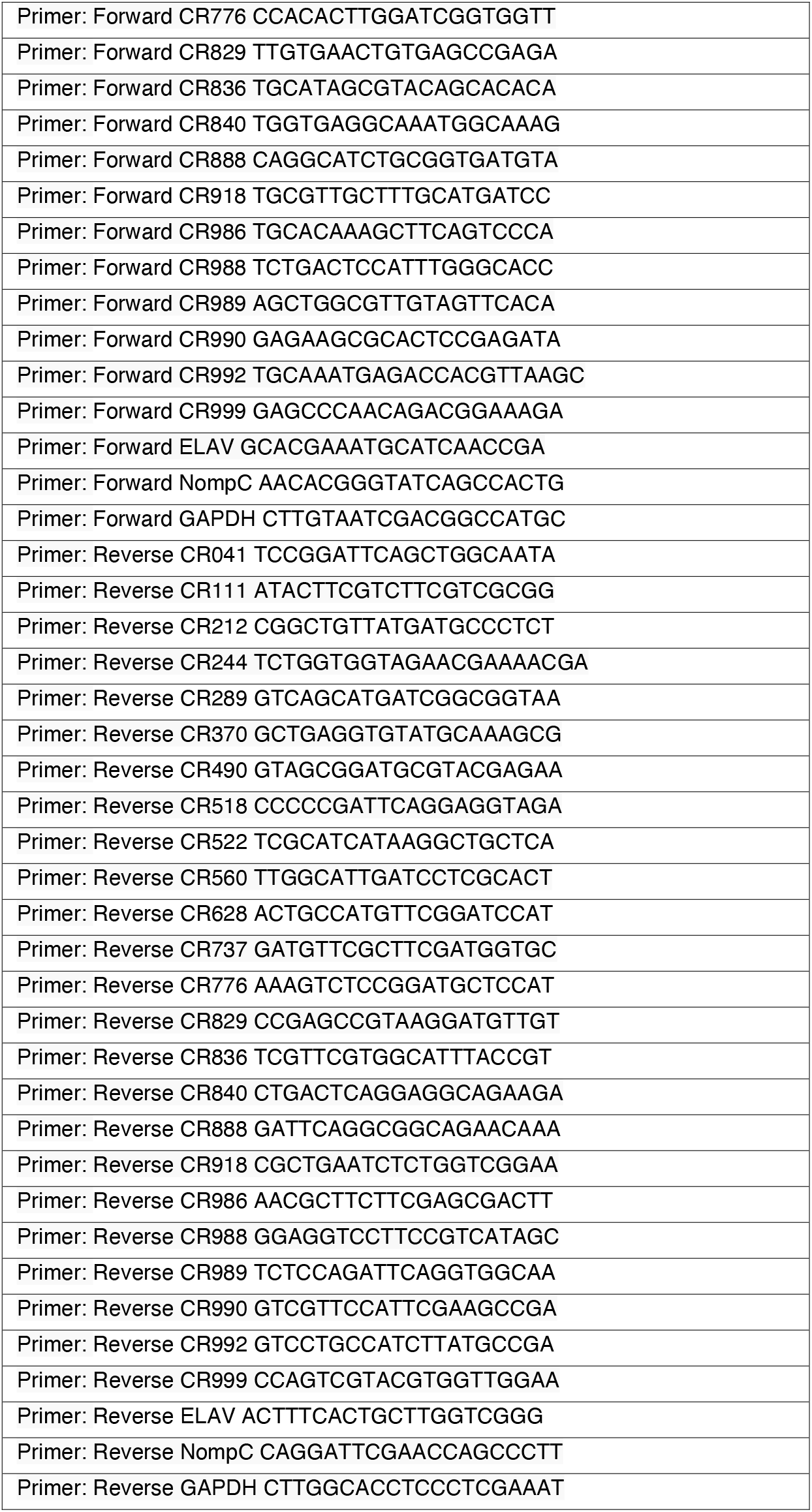
List with all oligonucleotide DNA primers used for quantitative RT-PCR amplification. Primers were designed using the Primer-BLAST tool from NCBI and validated by *in silico* PCR on the reference *O. bimaculoides* genome.

**Table S5.**
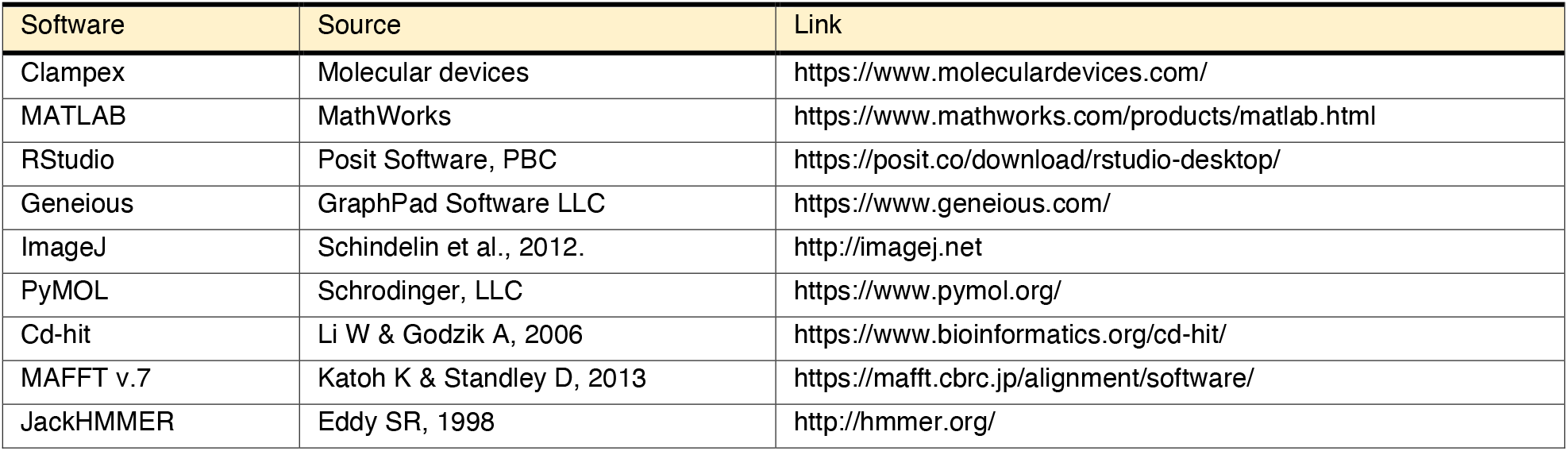
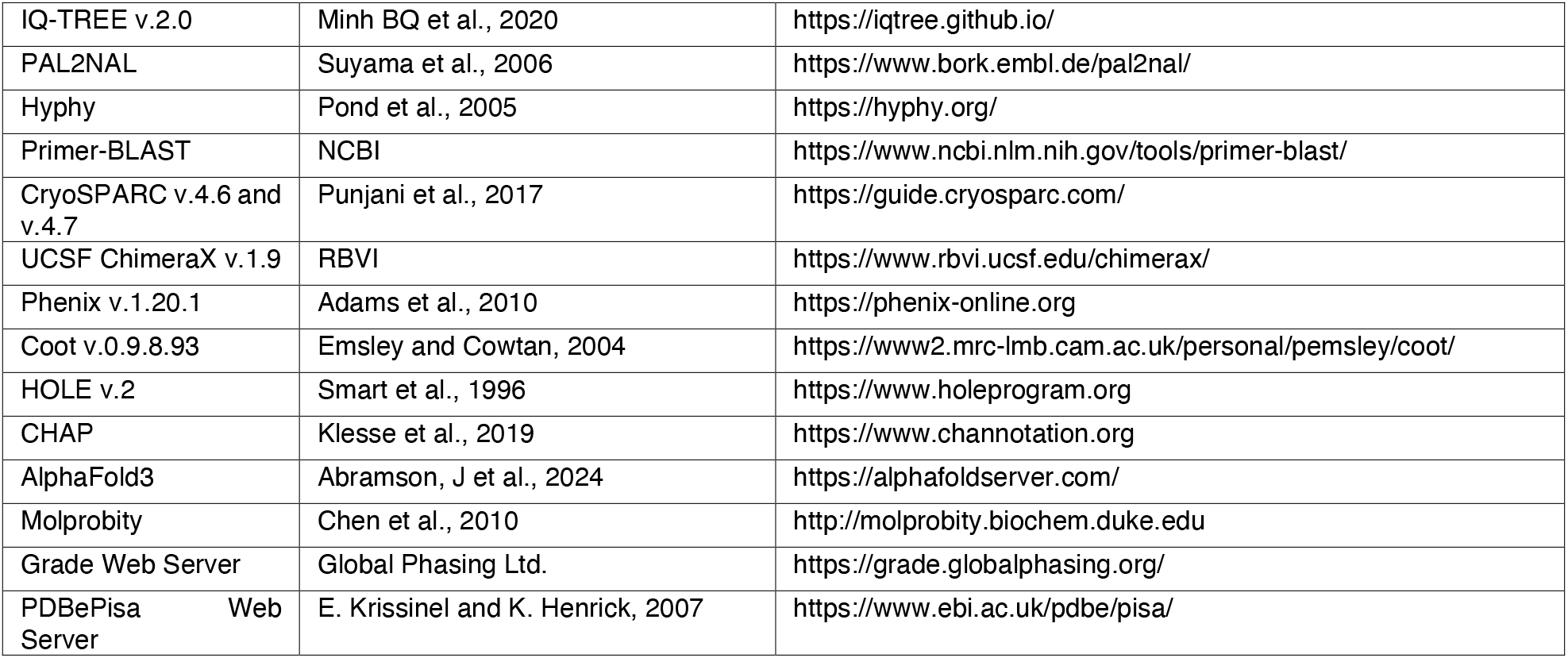
List with software used for bioinformatic and data analyses and their source.

**Table S6.**
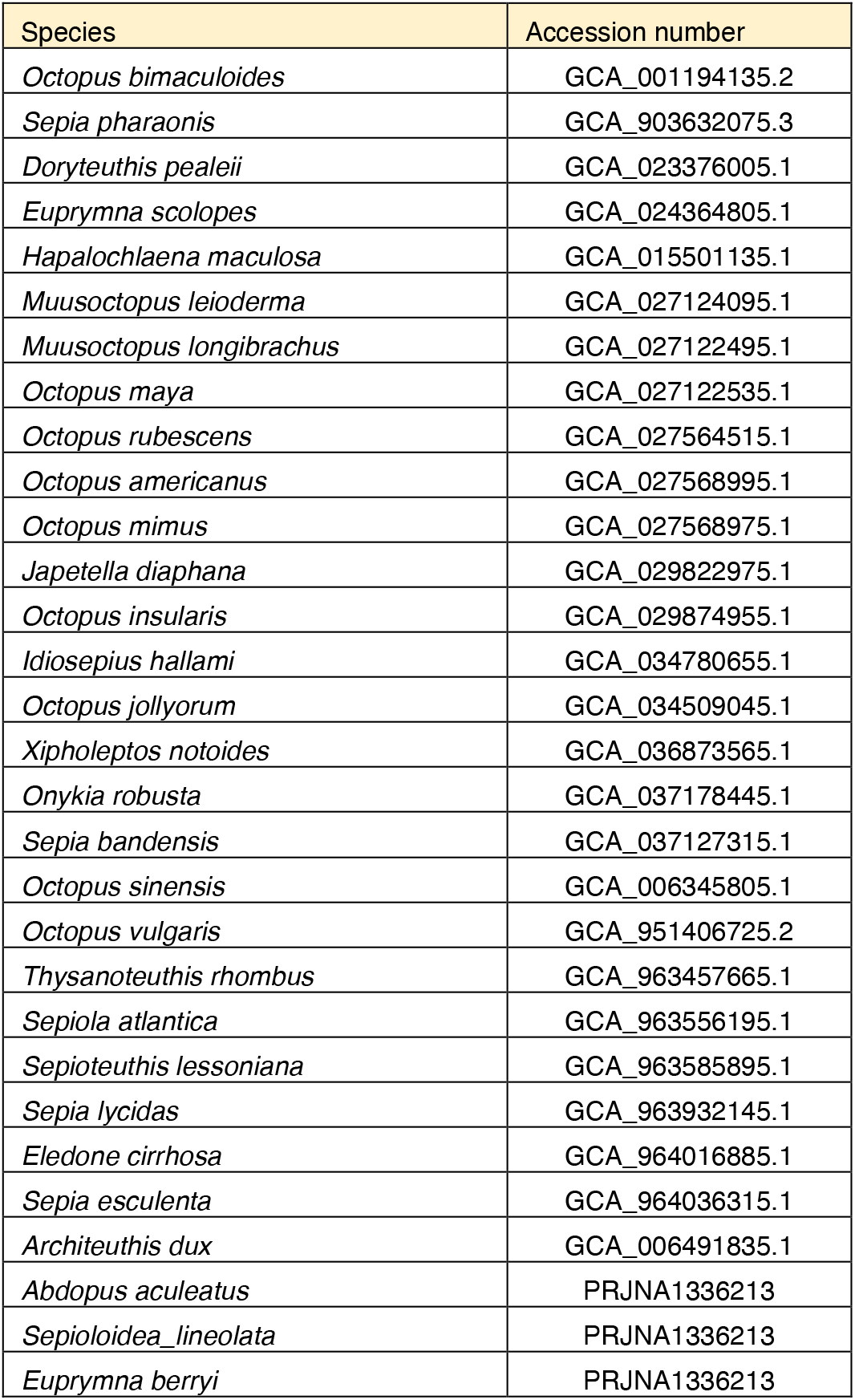
Cephalopod species and GenBank RefSeq or NCBI Sequence Read Archive accession numbers used to mine *CR* and *AChR* like genes.

**Table S7.**
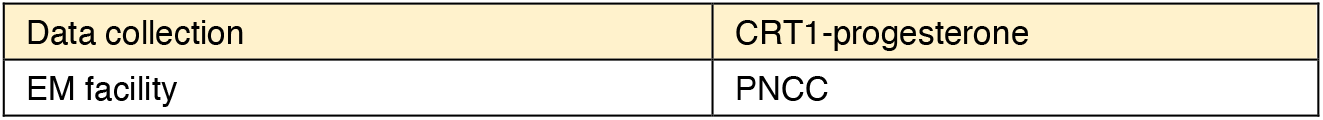
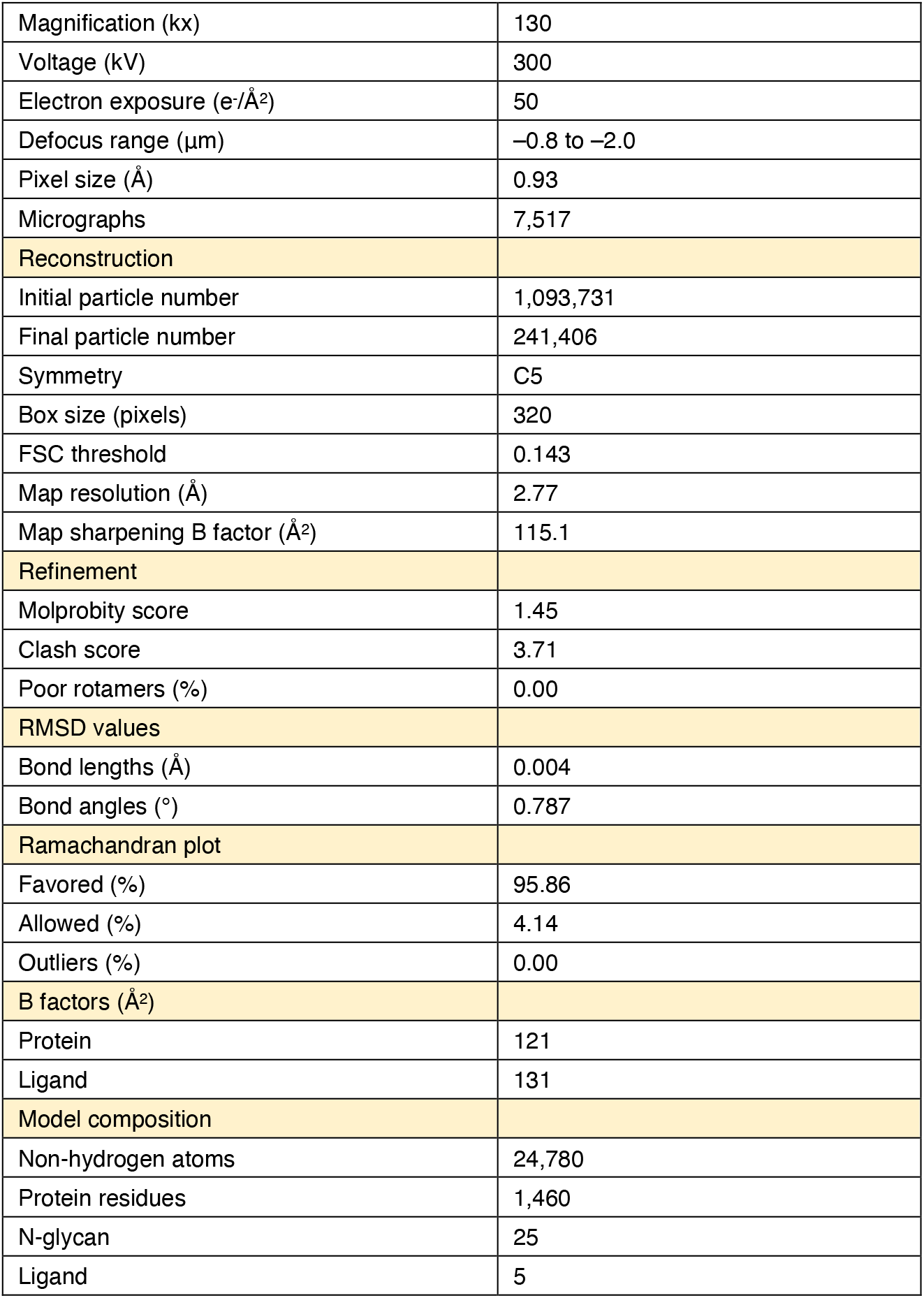
Cryo-EM data collection and refinement statistics for CRT1-progesterone complex shown in **Fig. 4.**

|  |  |
| --- | --- |
| Data collection | CRT1-progesterone |
| EM facility | PNCC |
| Magnification (kx) | 130 |
| Voltage (kV) | 300 |
| Electron exposure (e-/Å<sup>2</sup>) | 50 |
| Defocus range (μm) | −0.8 to −2.0 |
| Pixel size (Å) | 0.93 |
| Micrographs | 7,517 |
| Reconstruction |  |
| Initial particle number | 1,093,731 |
| Final particle number | 241,406 |
| Symmetry | C5 |
| Box size (pixels) | 320 |
| FSC threshold | 0.143 |
| Map resolution (Å) | 2.77 |
| Map sharpening B factor (Å<sup>2</sup>) | 115.1 |
| Refinement |  |
| Molprobity score | 1.45 |
| Clash score | 3.71 |
| Poor rotamers (%) | 0.00 |
| RMSD values |  |
| Bond lengths (Å) | 0.004 |
| Bond angles (°) | 0.787 |
| Ramachandran plot |  |
| Favored (%) | 95.86 |
| Allowed (%) | 4.14 |
| Outliers (%) | 0.00 |
| B factors (Å<sup>2</sup>) |  |
| Protein | 121 |
| Ligand | 131 |
| Model composition |  |
| Non-hydrogen atoms | 24,780 |
| Protein residues | 1,460 |
| N-glycan | 25 |
| Ligand | 5 |

**Movie S1. Octopus mating**. In this representative video two wild-caught *Octopus bimaculoides* were placed in a tank separated by an opaque barrier with small openings through which they could interact. After few minutes, males introduced their hectocotylus and maneuvered towards the female mantle and ovary. During this time both animals stopped all movements and locked in this position for around one hour after which they voluntarily separated. 1X speed.

**Movie S2. Tube assay**. In this representative video, a progesterone coated tube was presented together with an aversive CR agonist (nootkatone) or vehicle control (DMSO). Males expecting to mate moved towards the wall and extended their arms to investigate the other side. Male octopuses preferentially explored tubes coated in progesterone compared to other steroids, bile acids, terpenes, bitter molecules or vehicle control. 1X speed.

